# Decoding hidden goal-directed navigational states and their neuronal representations using a novel labyrinth paradigm and probabilistic modeling framework

**DOI:** 10.1101/2025.11.13.688348

**Authors:** Patrick S. Honma, Shreya C. Bangera, Reuben Thomas, Nicholas Kaliss, Dan Xia, Jorge J. Palop

**Affiliations:** Gladstone Institute of Neurological Disease, San Francisco, CA, 94158, USA; Neuroscience Graduate Program, UCSF, San Francisco, California, USA; Department of Neurology, University of California, San Francisco, CA, 94158, USA; Denali Therapeutics, South San Francisco, CA

## Abstract

Goal-directed navigation involves a sequence of planned actions aimed at achieving long-term goals through reinforcement, but detecting hidden states that support this process and their neuronal substrates remains a fundamental challenge. To address this, we developed a complex labyrinth test that mimics naturalistic foraging and implemented a novel hierarchical probabilistic modeling framework, Cognitive Mapping of Planned Actions with State Spaces (CoMPASS). This framework infers a nested state structure, comprising short-term surveillance–ambulation states (Level 1) and long-term goal-oriented navigational states (Level 2). Using CoMPASS, we show that successful navigation in wild-type mice is marked by increased recruitment of both surveillance and goal-oriented states specifically at decision nodes, revealing how sequential behavioral decisions culminate in long-term goals. In contrast, the humanized *App*^SAA^ mouse model of Alzheimer’s disease (AD) exhibited navigational impairments marked by diminished surveillance during decisions, reduced goal-directed states, and increased navigation stochasticity. Importantly, we show that gamma oscillations in the posterior parietal cortex (PPC), a region involved in spatial navigation planning, encode these CoMPASS behavioral states and their dynamic operating modes linking spatial locations to long-term goals. Our findings provide a novel paradigm for assessing hidden goal-directed navigational states and identify gamma oscillations in the PPC as their neural substrates.

## Introduction

Goal-directed behavior is a latent state that involves planned actions directed toward achieving specific objectives and is foundational for adaptive, value-based cognitive processes. Unlike non-goal-directed or spontaneous behavior, goal-oriented behavior incorporates feedback and reinforcement to optimize performance over time. Foraging—an evolutionarily conserved behavior—exemplifies this capacity, as animals navigating complex environments generate both spatial and value-based cognitive maps^1,2^ of optimal routes through surveillance and reinforcement and update strategies based on internal value-based states beyond immediate sensory and spatial cues. Although significant progress has been made identifying spatial map representations^3–5^, detecting hidden behavioral states of surveillance and goal-directed planning remains challenging, as these latent cognitive processes can only be defined through computational modeling during rewarded navigation tasks. Developing new approaches for detecting hidden states of reinforced learning is critical for elucidating the mechanisms underlying these cognitive processes and defining their neuronal representations in health and disease.

The neural mechanisms that support spatial representations and planning of goal-directed navigation likely involve coordinated neuronal activity across distributed networks. The posterior parietal cortex (PPC) represents a promising candidate, as PPC neurons track internal spatial representations of navigational routes^6^, encode error-correction signals^7^, monitor self-motion through space^6^, and engage in prospective movement coding^8^. Unlike the hippocampus, which supports allocentric spatial representations, the PPC integrates sensorimotor information with planned routes and choice-specific trajectories during navigation^9^, potentially linking spatial representations to varying long-term navigational goals^10^. Interestingly, lesions of the posterior parietal cortex (PPC) in humans lead to deficits in spatial orientation, particularly in tracking navigational routes^11^. PPC dysfunction has also been implicated in Alzheimer’s disease (AD)^12–17^, where early impairments in spatial planning and navigational flexibility may reflect disruptions in PPC-mediated integration of internal goals with spatial information.

Reinforcement learning (RL) computational modeling has been instrumental in formalizing how animals assign value to behavioral outcomes through reward prediction and feedback-based learning, linking decision-making behavior to neuronal dynamics such as dopaminergic prediction error signaling^18^ and hippocampal state-space encoding^19,20^. These models often discretize natural behavior into fixed action-outcome contingencies that do not capture the continuous, varying hidden states during sustained goal-oriented navigational tasks. Unsupervised modeling frameworks such as Hidden Markov Models (HMMs)—a class of state space models—have proven to excel at capturing behavioral variability and continuously evolving hidden behavioral states. They have been applied to infer latent states from coordinate location data of animals foraging in the wild^21,22^ or to identify structured trajectories in rodent navigation tasks^23^. However, traditional HMMs analyze only immediate behavioral patterns without incorporating how these states relate to broader goals or reward contexts. What remains unexplored is the use of state space models to characterize behavior as a temporally structured goal-directed process. Unlike the established use of state space models to extract latent structure from population-level neural activity^24,25^, there is no comparable framework for behavior that models it as a sequential process, where decisions at each step support long-term objectives.

To address these limitations, we developed a hierarchical probabilistic modeling framework— Cognitive Mapping of Planned Actions with State Spaces (CoMPASS)—that decomposes behavior into two interconnected levels: Level 1 infers short-term surveillance-ambulation behavioral states through kinematic patterns, while Level 2 assesses long-term goal-oriented navigational states through spatial and value-based metrics relative to a target. Unlike prior approaches, CoMPASS treats behavior itself as a dynamic, evolving state space, demonstrating how behavioral trajectories alone carry sufficient structure to reveal internal planning dynamics. We paired this computational framework with a complex labyrinth maze designed to promote rewarded goal-directed navigation and conducted electrophysiological recordings to interrogate the neural substrates of these hidden behavioral states. We identified well-defined latent navigational states, including surveillance and goal-directed behaviors, that distinguished successful from unsuccessful navigation. Remarkably, gamma oscillations in the PPC differentiated key elements of goal-directed navigation captured in our CoMPASS states, including motor planning and binding of spatial context with long-term reward information, and were predictive of successful navigation. This tight link to underlying neuronal activity both validates and strengthens our behavioral modeling results. Furthermore, the *App*^SAA^ mouse model of AD displayed deficits in hidden states associated with impaired navigational performance. Together, these findings establish behavior-centered state space modeling as a powerful approach for revealing latent goal-directed navigation processes, identifying PPC gamma oscillations as a critical brain region and neuronal mechanism of navigational planning.

## Results

### A rewarded multi-path navigational task to assess latent behavioral states of spatial navigation

We developed a complex labyrinth maze to assess hidden behavioral states during spatial learning. The labyrinth maze consists of a 12 x 12 grid (144 nodes) that contains 24 decision-making nodes (22 three-way, 2 four-way), 14 non-reward zones (4 loops, 10 dead ends), and a reward zone (target) with sucrose pellets (**Fig. 1a**). The maze features an optimal route (reward path) to the target zone. The probability of a successful bout (home cage to target) is 4.5x10^-6^, which is substantially lower than that of the most complex mazes, including the Cincinnati (1.56x10^-2^)^31^, Barnes (5.0x10^-11^)^32^ mazes, and the Rosenberg labyrinth maze (1.6x10^-2^)^15^. Mice are free to navigate between their home cage and the target zone during a single, 13-hour overnight trial in dark conditions. Behavior is recorded with near-infrared cameras from a ventral view, and analyzed with DeepLabCut^15,16^ to track body landmarks for pose estimation (**Fig. 1b**). Wild-type (WT) mice initially explored all maze zones (**Fig. 1c**, blue), but over time they displayed a progressive increase in reward path preference and a decrease in exploration of non-reward zones, including loops and dead ends (**Fig. 1c**, yellow), indicative of learning. Quantification of each exploratory bout—defined as a single, uninterrupted exploration into the maze and back to the home cage—revealed a rapid decrease of deviation (time) from the reward path within the first ∼100 bouts that persisted through the trial (∼450 exploratory bouts) (**Fig. 1d**). Notably, outward bound trajectories (towards the target) showed less deviation from the reward path compared to inward bound trajectories (toward home) (**Supplementary Fig. 2a**), establishing the goal-oriented nature of navigation toward the target zone. Additionally, mouse speed increased with successive exploratory bouts (**Fig. 1d**), indicating continuous engagement with the task throughout the trial without detectable habituation.

**Figure 1.**
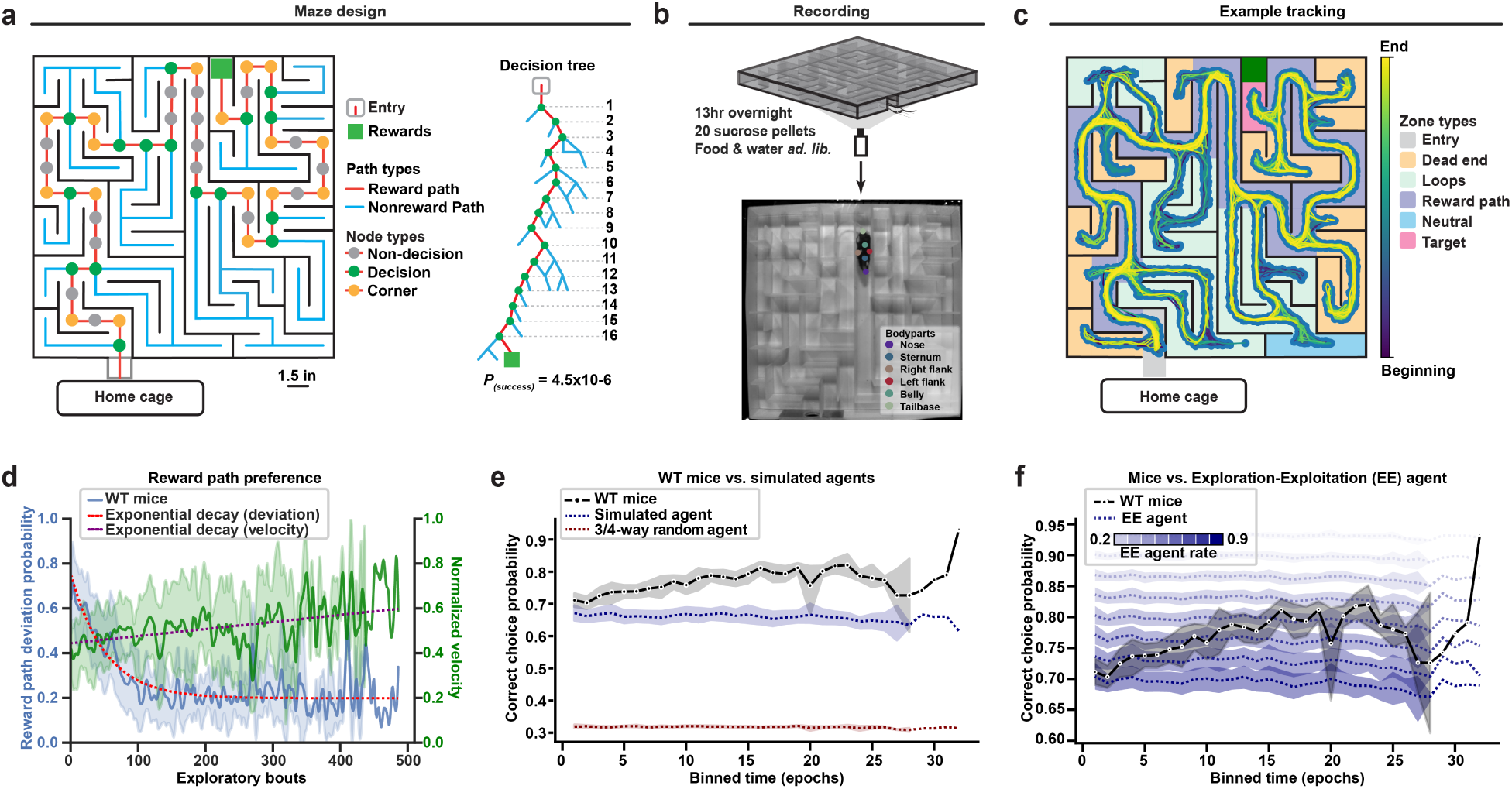
Development of a novel labyrinth maze for assessing latent behavioral states of spatial navigation. 3-7-month-old wildtype (*n*=17; 7F, 10M) mice were tested in the labyrinth maze**. a**, Design of the labyrinth maze with path and node types highlighted (left). The labyrinth maze decision tree showing the 16 decision nodes along the reward path to the target zone (right). **b**, Each trial consists of one, 13-hr overnight trial in which mice are free to navigate between their home cage and the labyrinth maze (top). Mouse keypoints are tracked using DeepLabCut, and the sternum allocentric coordinates are used for downstream analyses (bottom). **c**, Heatmap plot showing the mouse trajectories from the beginning (blue) to the end (yellow) of the trial (13 hours, overnight), where connected points indicate consecutive points in time. Grid nodes are colored by zone types. **d**, Deviation from the reward path from a cohort of WT mice, showing reduced deviation from the reward path with more exploratory bouts into the maze. Exploratory bouts are defined as any trajectory into the maze and back home, irrespective of reaching the target. Lines and shaded areas represent mean of mice ± sem. **e**, Probability of staying on the reward path at decision nodes, as compared to a simulated agent and a 3- or 4-way random agent. Probability is quantified during binned epochs of non-stationary transitions in the maze. **f**, Probability of staying on the reward path for WT mice compared to a simulated agent with varying exploration-exploitation (EE) rate. High EE rate represents higher exploration. Lines and shaded areas represent mean of agents ± bootstrapped CI (**e,f**).

To assess randomness in decision-making, we compared mouse navigation behavior against a simulated agent that replicates experimental mouse trajectories but chooses direction randomly at decision nodes (**Fig. 1e**) or an agent that operates with varying exploration-exploitation rates (**Fig. 1f**) during navigation. For this analysis, we defined epochs as discrete chunks of time considering transitions to unique grid nodes throughout the maze. These simulated agents made decisions based on empirically derived probabilities of correct choices—defined as continuing on the reward path at decision nodes. WT mice significantly outperformed the simulated random agent (exploration rate of 1.0; **Fig. 1e and Supplementary Fig. 2b,c**). Interestingly, mice progressively refined their navigation strategies with an initial higher exploration phase that gradually decreased as navigation became more exploitative and goal-directed (**Fig. 1f**). Mice outperformed the agents with 0.6–0.9 exploration rates, as evidenced by higher probability of staying on the reward path at decision nodes (**Fig. 1f**). We also tested a purely random agent using random probability choices (0.5 for binary, 0.33 for tertiary; **Fig. 1e**). As expected, performance with this agent was much lower than with both the simulated random agent and the exploration-exploitation agent, confirming that mice engage in non-random, goal-directed behavior. Overall, WT mice rapidly learned the maze’s goal-directed nature during a single overnight trial, showing increased exploitation behavior and preference for the reward path.

### Assessing spatial navigation in a complex labyrinth maze in WT and *App*^SAA^ mice

To assess spatial navigation in this newly developed complex labyrinth maze, we first generated heatmaps of zone preference and calculated the randomness of zone usage over time. Heatmap analysis of zone preference over time showed that mice rapidly developed target preference during the overnight trial (**Fig. 2a**). Although WT mice initially explored the target zone and a neutral zone equidistant from the home cage equally (**Fig. 2b**), they quickly learned to prefer the target zone, demonstrating progressive enhancement of reward-seeking behavior. This learning was accompanied by decreasing Shannon’s entropy values over time (**Fig. 2c**), indicating reduced behavioral stochasticity and increased targeted behavior. WT mice also showed significant path optimization, developing increasingly shorter paths to the target zone and longer paths to the neutral zone after repeated visits (**Fig. 2d,e**). The probability of successful bouts also increased over time (**Fig. 2f**) closely paralleled increases in target zone occupancy (**Fig. 2b**), confirming that WT mice not only reached targets more frequently but also developed coherent navigational patterns. We next tested whether this novel labyrinth maze detects navigational deficits in the humanized *App*^SAA^ knock-in model of AD^29^. *App*^SAA^ mice displayed reduced target zone preference (**Fig. 2b and Supplementary Fig. 4h-k**) and increased stochasticity (**Fig. 2c**). *App*^SAA^ mice also exhibited significantly reduced success rates with flatter learning curves (**Fig. 2f**) and showed impaired target search efficiency, requiring longer path lengths after 20 visits compared to the progressively shorter paths developed by WT mice (**Fig. 2d,e**). Altogether, WT mice demonstrated rapid and sustained overnight learning, efficiently navigating toward the target zone via the optimal path (reward path) while progressively reducing their randomness and increasing occupancy of goal-relevant target zones. Although *App*^SAA^ mice also learned (**Supplementary Fig. 3**), they exhibited clear navigational deficits characterized by delayed learning, persistent stochastic behavior, reduced reward-seeking preference, and impaired path optimization.

**Figure 2.**
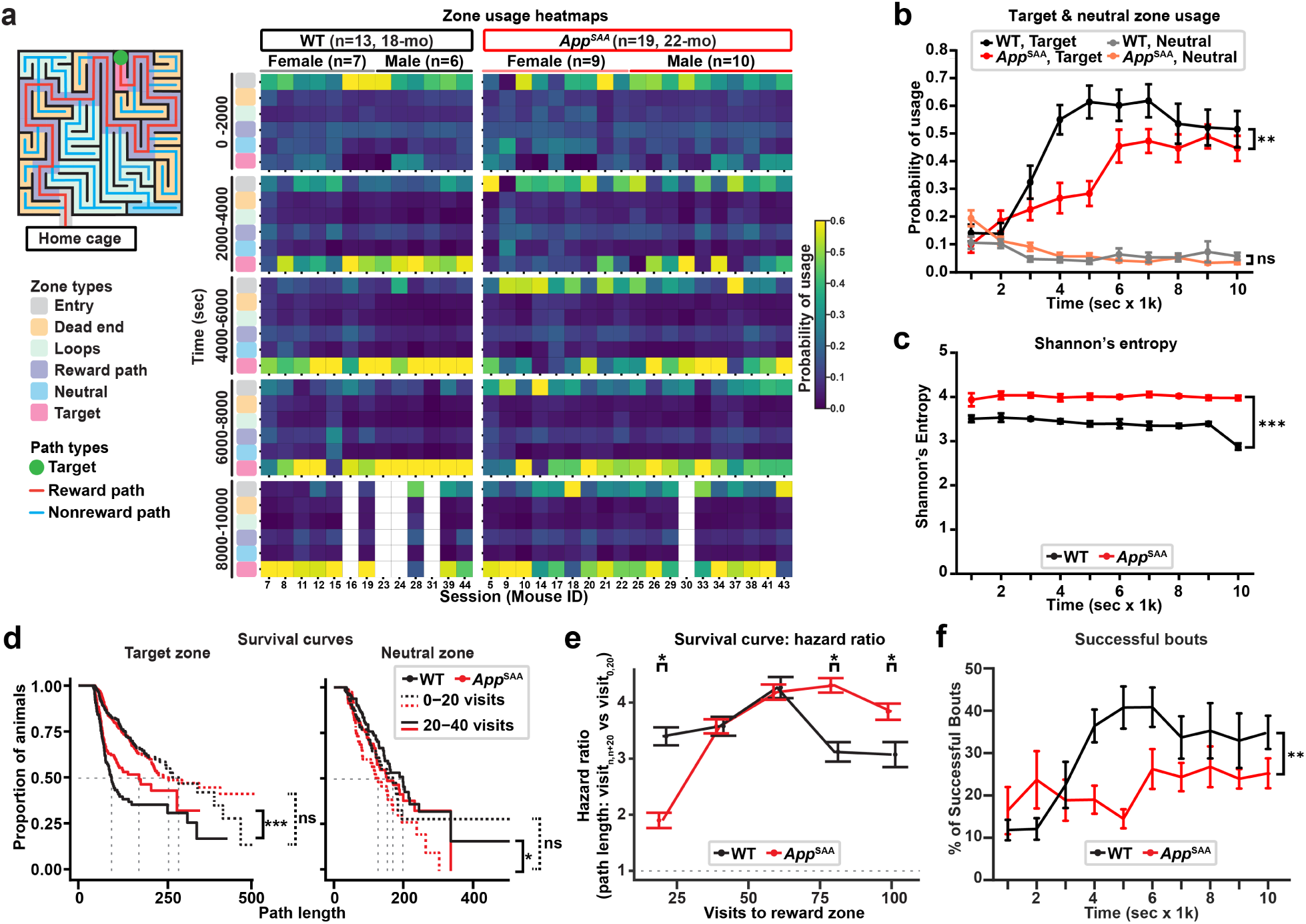
Wild-type mice develop directed target engagement while *App*^SAA^ mice show navigational stochasticity. 22-month-old *App*^SAA^ (*n*=19; 9F, 10M) and WT (*n*=13; 7F, 6M) mice were tested in the labyrinth. **a**, Labyrinth maze diagram indicating zone and path types (left). Heatmap showing the probability of usage of each labyrinth zone over time (right). **b,** Quantification of the zone usage heatmaps for the target zone and neutral zone. Lines and error bars represent mean ± sem. Repeated two-way ANOVA genotype x time interaction, \*\**p*<0.01. **c**, The randomness of exploration was estimated by the Shannon’s entropy of each bin of the heatmaps in (**a**). **d,** Kaplain-Meier survival curves showing *App*^SAA^ mice take longer path lengths to reach the target zone with 20-40 prior visits to the reward zone (left), and WT mice take significantly longer path lengths to visit the neutral zone with 20-40 prior visits to the neutral zone (right). Cox mixed effects model, \**p*<0.05, \*\*\**p*<0.001. **e,** Hazard ratio of the survival curves comparing the path lengths (**d**) during successive 20-visit bouts to the target zone compared to the path lengths during the first 20 visits. Mean ± 95% confidence intervals. Non-overlapping error bars indicate rejection of null hypothesis with \**p*<0.05. **f**, Quantification of successful bouts (home cage to target and back) to the target zone in each 500 sec bin. Lines and error bars represent mean ± sem. Repeated two-way ANOVA genotype x time interaction, \*\*\**p*<0.01. Lines and error bars represent mean ± sem (**b,c,f)**.

### Hidden behavioral states reflect successful navigation

To investigate hidden behavioral states underlying navigation, we leveraged the first level of our CoMPASS modeling framework, which identifies short-term behavioral states based on kinematic patterns using a discrete-time Hidden Markov Model (HMM) (**Supplementary Fig. 1**). We computed step length (distance traveled between frames) and turn angle (changes in heading direction) as inputs to the model (**Fig. 3a**). The momentuHMM^21^ package modeled these features and identified two latent behavioral states: “active surveillance” (short step length, high turn angle) and “automated ambulation” (long step length, low turn angle). As expected, surveillance was prominent at dead ends and corners, whereas ambulatory behavior occurred most frequently at the straight non-decision nodes (**Fig. 3b**). However, visits to decision nodes resulted in increased surveillance probability, consistent with behavioral state changes at these locations. Spatial mapping of state probabilities revealed genotype-specific differences, particularly at decision nodes along the reward path (**Fig. 3c**), indicating impaired hidden behavioral states in *App*^SAA^ mice. A quantification of surveillance state across all nodes and path types confirmed that the surveillance probability deficits were specific to decision nodes (**Fig. 3e**), indicating that *App*^SAA^ mice exhibit decision-and location-specific impairments in latent states engagement during navigation.

**Figure 3.**
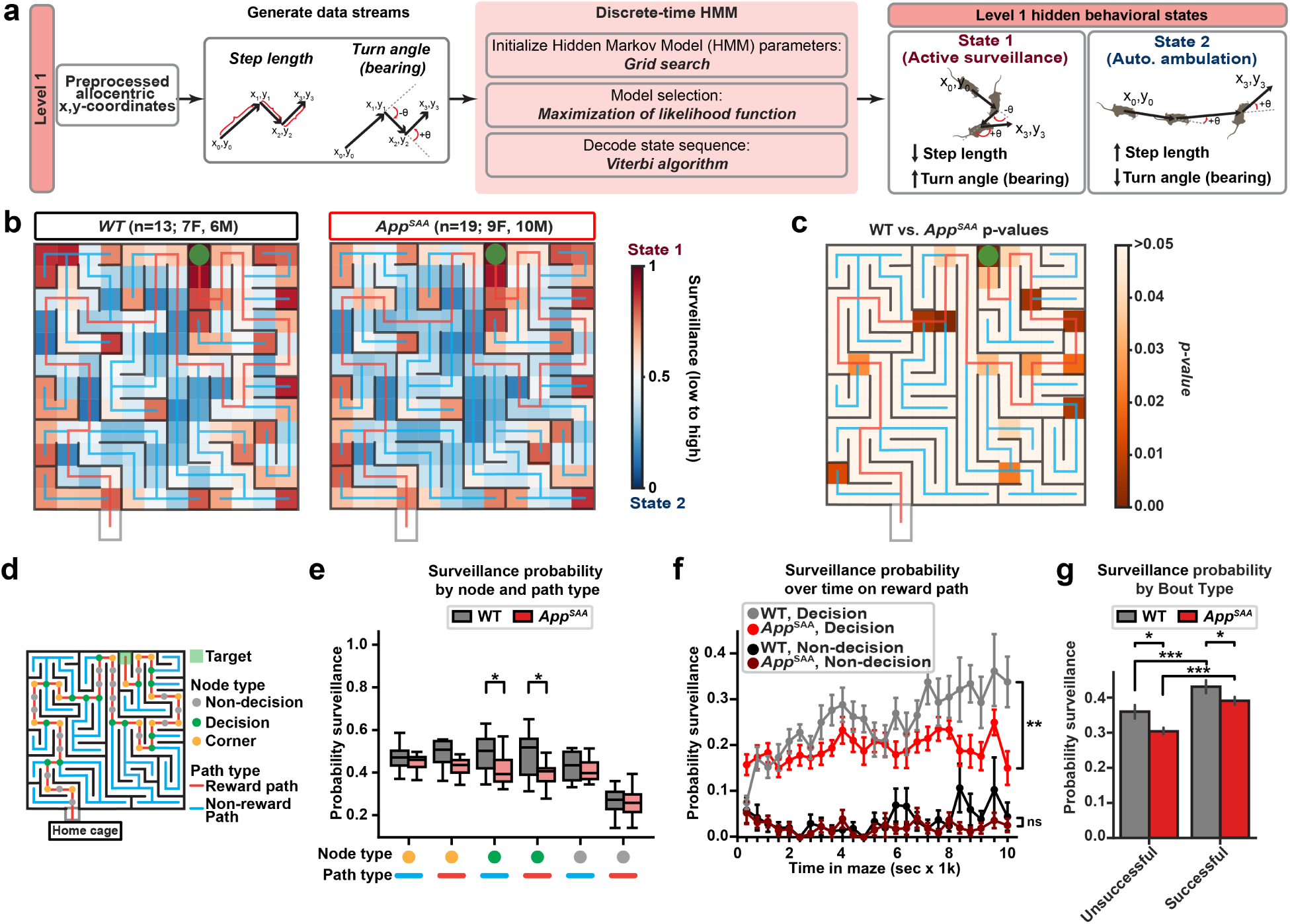
Deficits in hidden behavioral states at decision nodes as identified through discrete-time Hidden Markov Models. 22-month-old *App*^SAA^ (*n*=19; 9F, 10M) and WT (*n*=13; 7F, 6M) mice were tested in the labyrinth. **a**, The step length and turn angle were computed from the allocentric coordinates of the mouse and served as input for a two-state hidden Markov model (HMM). Two hidden behavioral states were identified, characterized by either low step length and high turn angles (active surveillance, red) or high step length and low turn angles (automated ambulation, blue). **b**, Heatmaps of the probability of being in surveillance (red) or ambulation (blue) in individual grid cells. **c**, Heatmap of the p-values comparing state probability between WT and *App*^SAA^ at each grid node. Independent t-test. **d**, Diagram showing node types (non-decision, decision, and corner) and path types (reward, non-reward). **e**, Boxplots showing the probability of active surveillance at the given node type and path type, showing reduced surveillance in *App*^SAA^ mice at decision nodes on the reward and non-reward path. Independent t-test. \**p*<0.05. **f**, Probability of active surveillance over time at decision and non-decision nodes along the reward path in 400 second bins, whereby *App*^SAA^ have reduced probability of being in the surveillance state at decision nodes. Lines and error bars represent median ± sem. Repeated two-way ANOVA for genotype x time interaction, \*\**p*<0.01. **g**, Mean active surveillance probability during successful and unsuccessful bouts for both genotypes. Bars and error bars represent mean ± sem. Linear Mixed Effects Model with mouse as a random factor, \**p*<0.05, \*\*\**p*<0.001.

To understand the nature of these differences, we analyzed the state probability at decision and non-decision nodes along the reward path over time. WT mice progressively increased their active surveillance probability specifically at decision, but not at non-decision, nodes along the reward path (**Fig. 3f**), indicating that enhanced surveillance at these critical navigational points is associated with successful learning (**Fig. 3f**). In contrast, *App*^SAA^ mice showed neither a progressive increase in surveillance nor comparable surveillance probability (**Fig. 3f**). Importantly, both genotypes showed heightened surveillance during successful bouts relative to non-successful bouts, indicating that surveillance state facilitates effective navigation (**Fig. 3g**). Our results suggest that our Level 1 CoMPASS states capture behavioral states linked to successful navigation in both spatial- and genotype-specific manners.

### PPC gamma oscillations reflect hidden behavioral states

The selective increases in surveillance state at the decision nodes in WT mice suggest contextualization of these critical spatial hubs during goal-directed navigation. To determine whether underlying neural activity distinguishes these latent states, we recorded wireless EEG from the posterior parietal cortex (PPC)–a region implicated in spatial trajectory planning^6^–in WT mice. Using multitaper power spectral density (PSD) analysis, we measured theta (5–12 Hz) and gamma (27–100 Hz) oscillations across spatial and temporal dimensions (**Fig. 4a**). Consistent with oscillations encoding speed^30^, mouse speed positively modulated the theta and gamma power (**Fig. 4b**). However, along the reward path, gamma power increased specifically at decision nodes relative to non-decision nodes (**Fig. 4c**, right), despite lower speed at these locations (**Fig. 4c**, left). Similarly, gamma power during the surveillance state was selectively enhanced at decision nodes along the reward path relative to non-decision nodes (**Fig. 4d**). This dissociation between gamma power and speed at decision nodes suggests that oscillatory activity in the PPC may encode hidden navigation states engaging in higher cognitive demand.

**Figure 4.**
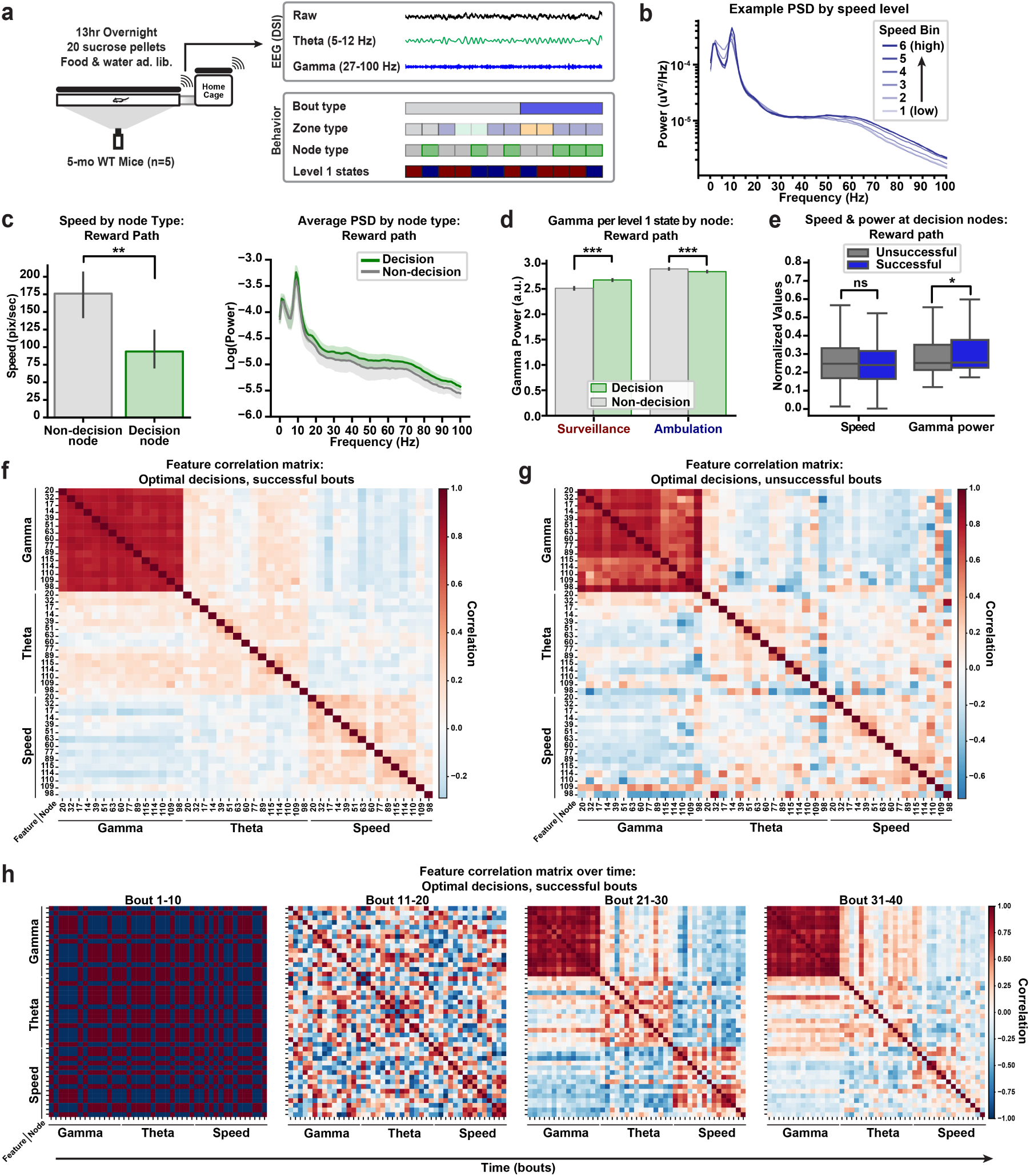
Neuronal synchronization in posterior parietal cortex (PPC) distinguishes successful navigation from spontaneous locomotor patterns. 5-mo-old WT mice (*n*=5, 5F) were implanted with wireless EEG/EMG probes in the PPC and recorded in the labyrinth maze during a 13 hr overnight trial. **A,** The power of theta and gamma oscillations were computed for each frame of the video and aligned to the behavioral results. **B**, Power spectral density (PSD) plots for different increasing speed bins in the maze (1–6; low to high speed), showing increased theta and gamma oscillations with increasing speed. **C**, Velocity for decision nodes and non-decision nodes along the reward path (left). Bars represent mean ± sem. Linear Mixed Effects Model with mouse as a random factor, \*\**p*<0.01. Median PSD corresponding by node type (right). Lines and shaded area represent median ± sem. **D**, Bar plots of gamma power for Level 1 CoMPASS states at decision nodes along the reward path and non-decision nodes along the reward path. Bars represent mean ± sem. Linear Mixed Effects Model with mouse as a random factor, \*\*\**p*<0.001. **e**, Boxplots showing the normalized speed and gamma power during successful and unsuccessful bouts. Linear Mixed Effects Model with mouse as a random factor, \**p*<0.05. **f,g**, Correlation matrix heatmap between gamma, theta, and speed at decision nodes during correct decisions (stay on reward path) during successful (**f**) and unsuccessful (**g**) bouts. Each row or column corresponds to a decision node. **H,** Feature correlation heatmaps as in (**f,g**) but in 10 bout bins.

To further test if gamma power encodes successful navigation, we assessed gamma power at decision nodes in successful and unsuccessful bouts. Consistently with the engagement of the PPC during navigation, gamma power was significantly higher during successful bouts than unsuccessful ones (**Fig. 4e**), with strong positive correlations across all decision nodes during correct decisions within successful bouts (**Fig. 4f**), indicating sustained and coordinated brain activity during successful navigation. However, this tight correlation was absent during correct decisions that ultimately led to unsuccessful bouts (**Fig. 4g**). Theta oscillations and movement speed showed much weaker relationships across decision nodes, indicating lower predictability of these variables encoding behavioral states (**Fig. 4f,g**). This stereotyped pattern of gamma power across decision nodes emerged progressively after approximately 20 bouts (**Fig. 4h**), indicating that these neural representations develop with successful navigational experience. Overall, these results offer a neural substrate for the behaviorally identified hidden states from Level 1 of our framework and for successful navigation. These findings are consistent with the role of individual neurons in the PPC in organizing planned actions and trajectories during navigation^6,8^.

### Detection of goal-oriented behavioral states using a multilevel HMM framework

While discrete-time HMMs are effective in characterizing short-term hidden behavioral states (Level 1), they do not fully capture how behavior is influenced by long-term goals and value-based navigation. Level 2 of our CoMPASS framework addresses this limitation by combining surveillance and ambulatory states (Level1) with four key aspects of goal-directed behaviors relative to the target location (Level 2): value-based distance (distance to target), oriented angle (immediate angle orientation to reward path), and Kernel Density Estimate (spatial distribution of node preferences) (**Fig. 5a**, *left*). This level employs a hierarchical Bayesian Gaussian Mixture Model (BGMM) and Gaussian Mixture Model Hidden Markov Model (GMM-HMM) architecture (See Supplementary Note) to cluster and model the combined data streams, revealing four latent behavioral states: oriented–surveillance, non-oriented–surveillance, oriented–ambulatory, and non-oriented–ambulatory (**Fig. 5a**, *right*).

**Figure 5.**
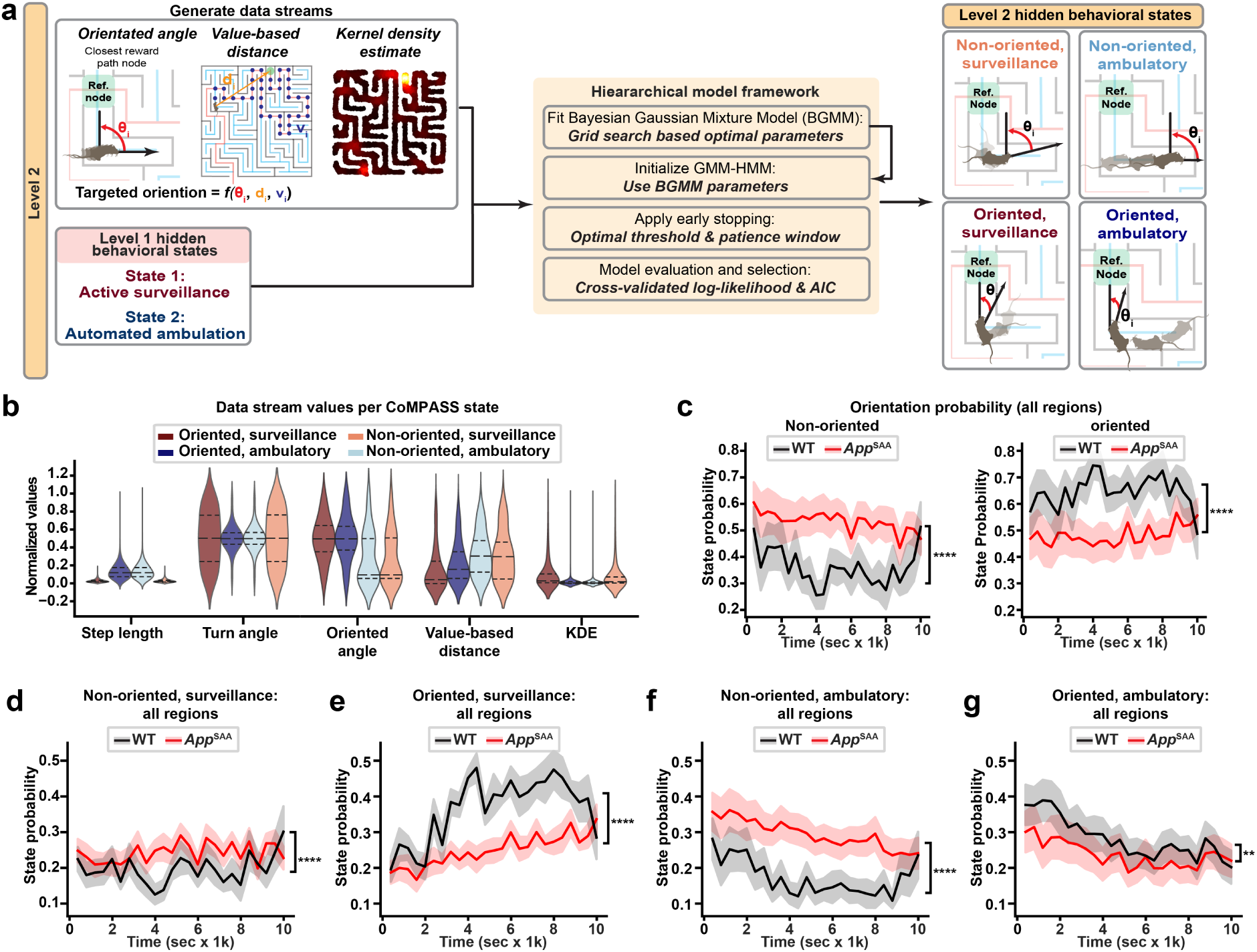
Level 2 CoMPASS states capture goal-oriented behavioral strategies that are disrupted in *App*^SAA^ mice. 22-month-old *App*^SAA^ (*n*=19; 9F, 10M) and WT (*n*=13; 7F, 6M) mice were tested in the labyrinth. **a**, Schematic of the hierarchical modeling framework to assess reward-guided behavioral states. The Level 1 hidden behavioral states along with Kernel density estimation of x-y coordinates, oriented angle, and value-based distance serve as data streams Level 2 of CoMPASS. Identified four hidden behavioral states: Embedded within the first level states (surveillance and ambulation) are the oriented or non-oriented states. **b,** Violin plots showing the distribution of data stream values per level 2 state showing increased sternum angle. **c**, Median probability of non-oriented (left) and oriented (right) states for WT and *App*^SAA^ mice. **d-g**, Median probability of (**d**) non-oriented-surveillance, (**e**) oriented-surveillance, (**f**) non-oriented-ambulatory, and (**g**) oriented-ambulatory over time. All statistics from repeated measures 2-way ANOVA for genotype factor, \**p*<0.05, \*\**p*<0.01, \*\*\**p*<0.001, \*\*\*\**p*<0.0001. Lines and shaded regions represent median ± sem.

Analyzing the distribution of data stream parameters confirmed clear separation among these four cohesive ethological behavioral states: ambulatory states showing greater step length, surveillance states showing greater turn angle dispersion, oriented states showing less oriented angle dispersion, non-oriented states showing higher value-based distances, and surveillance states showing increased KDE values (**Fig. 5b**). Random forest analysis across 50 runs revealed that all data streams contributed to state classification, with step length and oriented angle deviation emerging as particularly informative features (**Supplementary Fig. 5a**). Interestingly, WT mice exhibited higher levels of oriented states, including surveillance and ambulation, relative to corresponding non-oriented states across the full trial and all locations (**Fig. 5c-g**), indicating persistent goal-oriented behavior in WT mice. Conversely, *App*^SAA^ mice showed no preference between oriented and non-oriented states and had consistently higher probability of operating in non-oriented states (**Fig. 5d-g**). Relative to WT mice, *App*^SAA^ mice had also reduced probability of exhibiting oriented states and increased probability of non-oriented states, indicating disorientation in this goal-directed task (**Fig. 5c-g**). These differences in state probability were also apparent when analyzing behavior specifically on the reward path, with *App*^SAA^ mice showing the strongest deficits in oriented–surveillance state (**Supplementary Fig. 5c-g**). While WT mice spent most time in oriented states, *App*^SAA^ mice predominantly occupied non-oriented states (**Fig. 5c**). These robust genotype differences reveal behavioral alterations in goal-oriented navigational states that likely underlie task performance impairments. By bridging short-term behavioral patterns with long-term navigational goals, CoMPASS provides deeper insights into adaptive learning and spatial navigation processes.

### Gamma oscillatory dynamics in the PPC represent hidden navigation states

We next investigated whether short-term (surveillance) and long-term (goal-oriented) latent behavioral states reflect cortical neuronal synchronization in the PPC. Remarkably, goal-oriented states showed sustained and elevated gamma power compared to non-oriented counterparts at the reward path (**Fig. 6b**), decision nodes (**Fig. 6c**), and the rest of the maze (**Supplementary Fig. 7e**), suggesting spatial awareness of the orientation and likely increased cognitive demand during goal-directed navigation states. Consistent with Fig. 4, these oscillatory patterns were also dissociated from locomotor speed (**Supplementary Fig. 7b,d**), indicating that goal-directed navigation engages distinct synchronization mechanisms beyond simple kinematic patterns. Using Wasserstein Distance to compare gamma power distributions, we found that the oriented states, including oriented–surveillance and oriented–ambulatory, specifically increased gamma power along the reward path (**Fig. 6b**), particularly at decision nodes (**Fig. 6c**). These results indicate that oriented and non-oriented states represent distinct brain states, likely associated with distinct brain patterns of goal-directed and non-goal-directed behavior, respectively. To further test if dynamics of gamma power in the PPC encode short-term and long-term behavioral states, we assessed the temporal changes of gamma power in 8–second windows at the decision points. We found a specific increase in gamma power during oriented states, but a dip in gamma power during non-oriented states (**Fig. 6d**). Notably, a 3D UMAP representation of individual windowed gamma powers without behavioral state information revealed four distinct clusters corresponding to the four CoMPASS states (**Fig. 6e**), providing compelling evidence that the CoMPASS states represent distinct neural correlates for navigational states in the PPC.

**Figure 6.**
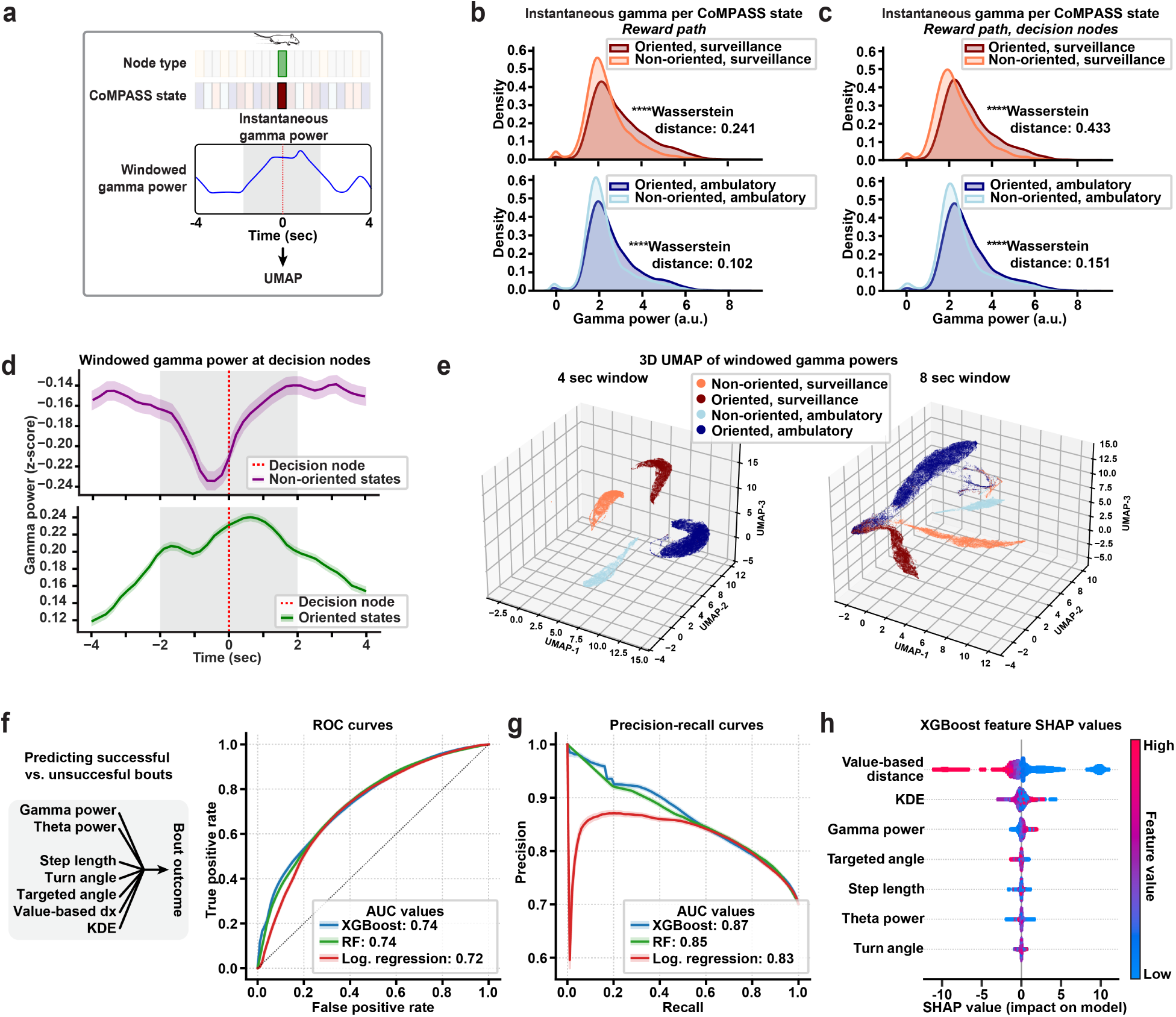
Gamma synchronization and hidden behavioral states distinguish goal-oriented from non-oriented states and predict successful navigation outcomes. 5-mo-old WT mice (*n*=5, 5F) were implanted with wireless EEG/EMG probes in the PPC and recorded in the labyrinth maze during a 13 hr overnight trial. **a**, Schematic showing calculation of instantaneous gamma (**b,c**) and 4- and 9-second-windowed of gamma power (**d,e**) when mice are at decision nodes along the reward path in specific CoMPASS states. **b,c**, Distribution of gamma power values in the four CoMPASS states along all nodes on the reward path (**b**) or specifically at decision nodes on the reward path (**c**). Wasserstein Distance Test, \*\*\**p*<0.001. **d,** Average z-scored gamma power around decision nodes when operating in non-oriented (top) or oriented (bottom) states. **e**, 3D UMAP of all windowed gamma powers colored by the CoMPASS states for 4 (left) and 8 sec (right) windows. Lines and shaded regions represent median ± sem. **f**, Classifying successful vs. unsuccessful bouts from neuronal features and all CoMPASS data streams using XGBoost, random forest, and logistic regression classifiers. ROC curves and the AUC values for the three classifiers. **g**, Precision-recall curves for the three classifiers in (**g**) and the AUC values. **h**, XGBoost feature importance plot with SHAP values for low (blue) to high (red) feature values.

Finally, we tested whether these neural signatures could predict navigational success. Using oscillation powers (theta and gamma) and the CoMPASS data streams to predict successful versus unsuccessful bouts (**Fig. 6f**), our models achieved high accuracy (**Fig. 6f**) and precision (**Fig. 6g**). As expected, close proximity to the target (low value-based distance) was the highest predictor of successful bouts. However, gamma power emerged as one of the highest predictors of successful bouts (**Fig. 6h**), establishing a critical role for gamma oscillations in PPC not merely as correlates of goal-oriented navigation states but potentially as a mechanism guiding effective spatial navigation in search of broader goals.

## Discussion

We developed CoMPASS, a hierarchical probabilistic modeling framework that identifies latent behavioral states underlying goal-directed spatial navigation. CoMPASS simultaneously infers short-term behavioral states (surveillance vs. ambulation) through instantaneous kinematic patterns and long-term states (goal-oriented states) through spatial and value-based metrics relative to the navigational target. We tested this computational framework in mice performing a novel complex labyrinth task that promotes goal-directed reward seeking and value-based spatial navigation. This approach revealed four well-defined latent navigational states that were dependent on both local and allocentric cues, and predicted successful navigation, suggesting that the defined states capture the spatial and value-based cognitive maps of spatial navigation. The neuronal underpinnings of these states were assessed by examining gamma oscillations in the PPC, which provided a neural signature that fully represented these latent behavioral states, with dynamics that evolved during learning and predicted successful navigation. Our computational framework and findings provide new insights into the neural mechanisms supporting goal-directed navigation and value-based spatial mapping, as well as their disruption in disease.

Goal-directed behavior involves the execution of planned actions to achieve specific objectives through dynamic value updating and strategy refinement. Behavioral sequences that lead to positive outcomes–such as reaching rewards–are reinforced over time, while the value of different actions is continuously updated based on accumulated experience. However, detecting these latent behavioral states remains challenging in part due to the difficulty of developing behavioral paradigms that engage continuous, long-term, reward-driven goal seeking (13 hr trials) using both spatial (location) and value-based cognitive maps, a dual requirement essential for modeling how animals integrate spatial representations with motivational states.

To meet this challenge, we developed a unique, complex labyrinth maze for eliciting goal-directed navigation in mice. Most traditional behavioral tasks (e.g., nose pokes, lever presses, T-mazes^31^, linear tracks, Barnes Maze^28^, labyrinth^32^) rely on binary or low-dimensional decision structures, short trials, and action spaces that cannot capture the continuous nature of real-world navigation. Our complex multi-nodal architecture integrates 24 decision points (3- or 4-way) along an optimal reward path, with a low probability of successful bout (4.5x10^-6^) (**Fig. 1a**), creating spatial uncertainty that demands continuous evaluation of exploration of various zones against exploitation of known reward paths. Both wild-type and *App*^SAA^ mice outperformed simulated random agents and exploration-exploitation agents at decision points (**Fig. 1e,f, Supplementary Fig. 2,3**), indicating structured strategies at key decision-making hubs. Moreover, mice dynamically shifted between exploration and exploitation based on spatial context and accumulated experience, suggesting that goal-directed navigation emerges from a mixture of instantaneous behavioral strategies embedded within the continuous series of learned planned actions.

This motivated the need for a computational framework capable of detecting temporally localized shifts in behavior while still inferring latent cognitive structure. While RL principles suggest that animals assign values to actions and locations based on experience^33^, our continuous paradigm with fixed reward contingencies makes traditional RL parameter estimation challenging. Rather than focusing on cumulative learning dynamics, we aimed to capture moment-to-moment behavioral states that reflect underlying cognitive processes supporting goal-directed navigation. HMMs excel at identifying latent behavioral states that evolve over time. While prior applications have been used to infer foraging or locomotor states from wild animal telemetry data^34,35^, newer implementations have successfully decoded distinct navigational strategies in spatial laboratory tasks^36^, providing a natural framework for inferring instantaneous behavioral states. Bayesian actor models formalize how goal-directed navigation can be modeled as optimal feedback control under uncertainty^37^—agents update estimations of spatial position, goals, and motor commands to dynamically guide planned actions towards long-term objectives. However, while flat HMMs effectively identify low-level patterns, they do not account for higher-order cognitive structure or the nested nature of decision-making inherent to goal-directed navigation.

CoMPASS thus draws on principles from RL, HMMs, and Bayesian actor models of navigation under uncertainty. Unlike RL—which specifies normative behavior given a reward model— CoMPASS inverts the inference: it uncovers latent structure directly from observed behavior, without assuming an optimal policy or reward function. This inversion is critical in naturalistic settings where agent goals, and cognitive maps must be inferred rather than imposed. CoMPASS leverages these theoretical principles to create data streams that encapsulate the continuous nature of our task: kinematic features that reflect immediate motor commands, spatial features capture goal-directed planning, and their integration reveals how animals alter short-term behavioral patterns in search of long-term goals. This hierarchical structure operates across multiple temporal and cognitive scales, capturing hidden states that reflect both short-term movement patterns (Level 1) and broader, goal-directed navigational objectives (Level 2).

Level 1 of CoMPASS employs discrete-time Hidden Markov models (HMMs) to infer short-term behavioral states from kinematic parameters (step length and turn angle), revealing two locomotor states: active surveillance and automated ambulation (**Fig. 3**). The surveillance state, characterized by slower movement and increased angular changes, resembles the deliberative processing underlying vicarious trial-and-error (VTE), where animals engage in orienting movements while mentally simulating potential trajectories and their associated rewards^38,39^. This behavioral signature of deliberative decision-making reflects distinct cognitive processes rooted in the computational demands of goal-directed navigation—the need to form an internal representation of the environment, estimate position relative to goals, and anticipate optimal routes to goal locations^40^. Critically, surveillance probability increased during successful bouts in both genotypes and was disrupted in *App*^SAA^ mice (**Fig. 3**), demonstrating that these kinematic states reflect underlying cognitive operating modes.

Level 2 extends this analysis to decode instantaneous goal-oriented behavior by incorporating spatial and value-based features: distance to target, oriented angle deviation from the optimal trajectory, and spatial distribution of events via kernel density estimation. Unlike tasks where animals can use direct vector-based navigation toward a visible target, our paradigm requires mice to engage in a series of iterative reorientation strategies in search of the target. This distinction is crucial—reward orientation cannot be captured by simple directional vectors but requires understanding the dynamic process of sequential reorientations that guide animals onto the reward path while maintaining the target as the ultimate objective. Level 2 proved remarkably sensitive to detect navigational impairments that were invisible to overt learning metrics, revealing that *App*^SAA^ mice showed the strongest deficits in oriented, surveillance state (**Fig. 5**)—a critical combination of deliberative processing and goal-directed orientation essential for successful navigation. While WT mice spent a majority of their time in oriented states, *App*^SAA^ mice predominantly occupied non-oriented states (**Fig. 5g**), indicating fundamental alterations in goal-directed strategy. This suggests that the increased stochasticity and impaired reward path optimization observed in *App*^SAA^ mice (**Fig. 2c,d**) stem from impaired ability to maintain goal-directed orientation during deliberative processing. By leveraging unsupervised clustering and temporal sequence modeling, CoMPASS offers an interpretable framework for characterizing the internal cognitive dynamics and their disruption in disease.

In search of the neural representations underlying our hidden states of goal-directed navigation, we recorded wireless EEG oscillatory activity from the PPC, which plays a critical role in spatial navigation through integration of spatial locations and route planning^6,41^. PPC neurons track animal’s self-motion and egocentric location of moving targets, with activity dynamically modulated by task demands^42^ and tuned to past, present, and future trajectories, enabling spatial-to-motor transformations for movement planning^2,8,10^. Consistent with these findings and the role of oscillations in spatial navigation^9–15^, we discovered that gamma oscillations in the PPC provided a well-defined neural representation of the four CoMPASS behavioral states (**Fig. 6e**). Most remarkably, each state showed distinct clusters of gamma oscillatory power dynamics around decision nodes (**Fig. 6e**), with particular increases when mice operated in goal-directed states that were reversed during non-goal-directed states (**Fig. 6d**). This suggests that PPC oscillations encode the individual components of goal-directed navigation captured by our framework, including motor commands, movement through space, planned trajectories, and deliberative processing.

While hippocampal theta sweeps and nested gamma oscillations are well documented in encoding spatial sequences and anticipated trajectories^38,43–48^, our results suggest that PPC could integrate reward-relevant spatial information into prospective goal-directed actions. In support of this, gamma power was not only highly stereotyped across all decision nodes on the reward path during successful bouts but also significantly increased during successful versus unsuccessful bouts (**Fig. 4g,h**) and emerged as a highly predictive feature of navigational success (**Fig. 6g-i**). This pattern suggests that the PPC may explicitly encode goal information, allowing for the transformation of allocentric spatial maps in the entorhinal cortex and hippocampus into body-centered representations that incorporate reward context. Our data demonstrates that exploiting the reward path versus exploring alternative trajectories produce distinct gamma representations in PPC, extending its role beyond spatial-to-motor transformations to include integration of spatial information with reward signals essential for successful goal-directed navigation.

While our findings demonstrate that PPC gamma oscillations provide a robust neural signature of our CoMPASS behavioral states, an important limitation is the lack of simultaneous hippocampal recordings. Given the critical role of the hippocampus in prospective trajectory planning^38,49^, it is possible that these representations are found in the hippocampus or other brain regions involved in goal-directed navigation. We anticipate that CoMPASS could have far-reaching impacts for decoding instantaneous behavioral states across a broad range of goal-directed navigational tasks, advancing our understanding of the mechanisms supporting this fundamental behavior.

## Supplementary Figures

**Supplementary Figure 1.**
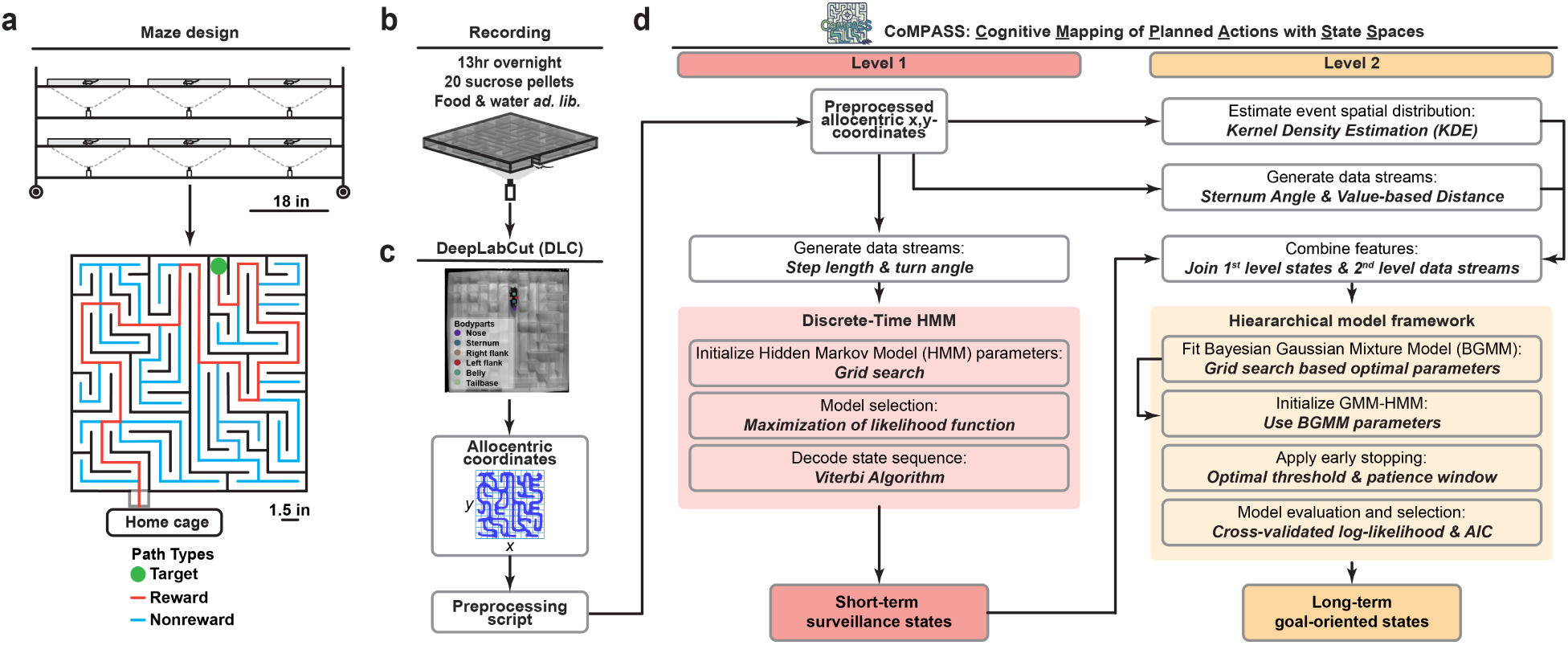
Labyrinth Design and Workflow. **a**, The design of our 6-labyrinth setup (top) and the design of each individual labyrinth maze (bottom). **b**, Each trial consists of one, 13-hr overnight trial in which mice are free to navigate between their home cage and the labyrinth maze. **c**, Mouse keypoints (nose, left flank, right flank, sternum, belly, tailbase) are tracked using DeepLabCut, and the sternum allocentric coordinates are used for downstream analyses. **d**, CoMPASS: A hierarchical modeling framework for uncovering hidden behavioral states that underlie spatial navigational strategies. Level 1 of the hierarchical framework uses step length and turn angle to identify basic locomotor hidden behavioral states. Level 2 uses the level 1 decoded states, a oriented angle, value-based distance, and kernel density estimation to uncover goal-directed hidden behavioral states.

**Supplementary Figure 2.**
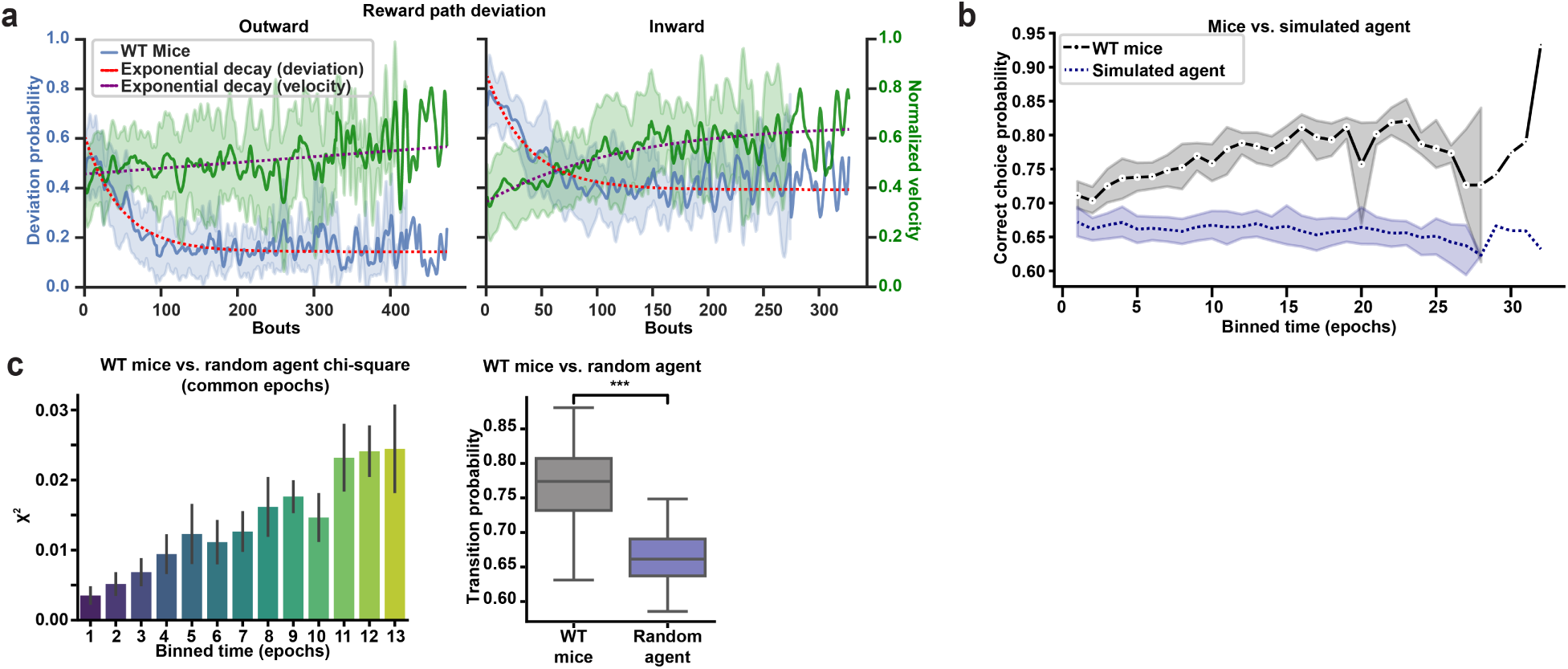
Validation of learning in WT Mice. 3-7-month-old wildtype (*n*=17; 7F, 10M) mice were tested in the labyrinth maze**. a**, Deviation from the reward path from a cohort of 3–7-month-old WT mice (*n*=17; 7F, 10M), showing reduced deviation from the reward path during outward (towards maze) and homeward (towards home) segments of bouts. Velocity increases with bouts for both cases. Lines and shaded area represent the mean ± sem**. b**, Probability of staying on the reward path at decision nodes, as compared to a simulated agent. Lines and shaded area represent the mean ± bootstrapped CI**. c**, The chi-square test for independence between WT mice and the random agent for optimal transitions at decision nodes, only considering a number of epochs in which all mice were performing (left). Bars represent mean ± sem. Boxplots showing probability of optimal decisions at decision nodes for WT mice and the simulated agent (right). Mann Whitney U-Test, \*\*\**p*<0.001.

**Supplementary Figure 3.**
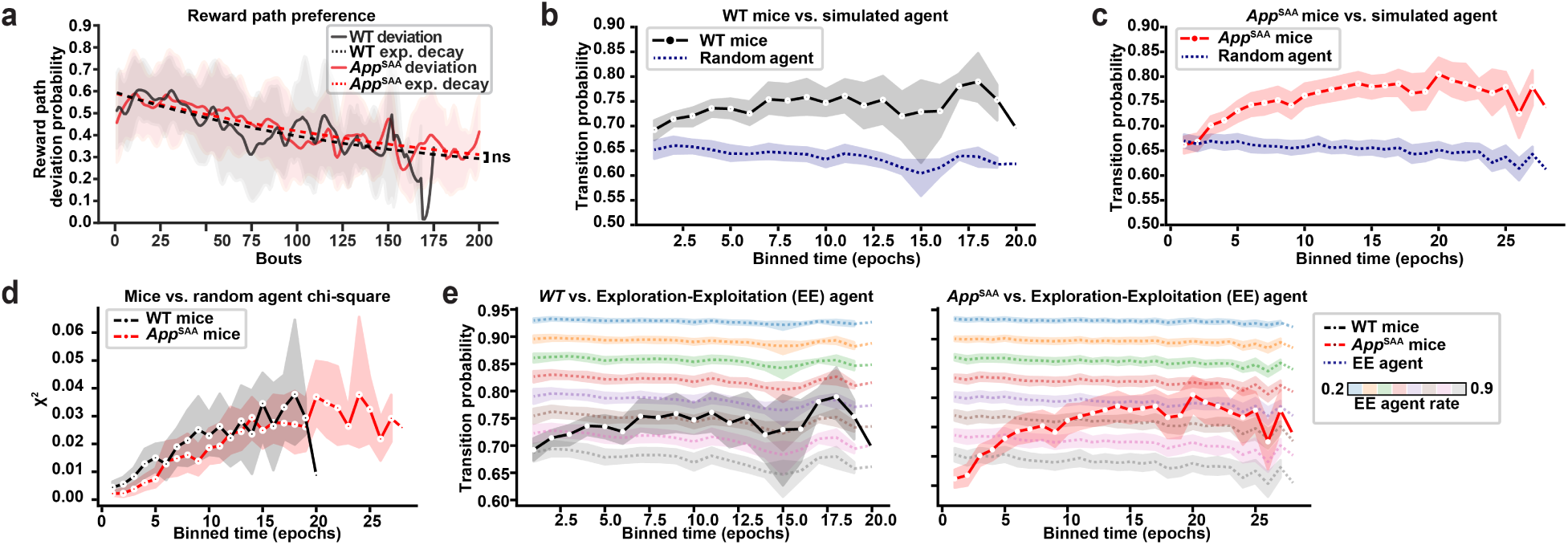
Validation of learning and performance in WT and *App*^SAA^. 22-month-old *App*^SAA^ (*n*=19; 9F, 10M) and WT (*n*=13; 7F, 6M) mice were tested in the labyrinth. **a**, Deviation from the reward path from *App*^SAA^ and WT mice, showing no differences in deviation probability. Median ± sem are shown. **b-c**, Probability of staying on the reward path at decision nodes for WT and *App*^SAA^ mice, as compared to a simulated agent. **c**, The chi-square test for independence between WT and *App*^SAA^ mice and the simulated agent for optimal transitions at decision nodes. **e**, WT (left) and *App*^SAA^ (right) mice compared to a simulated agent with increasing exploration-exploitation (EE) rate. High EE rate represents higher exploration. Lines and shaded area represent mean ± bootstrapped CI (**b-e**).

**Supplementary Figure 4.**
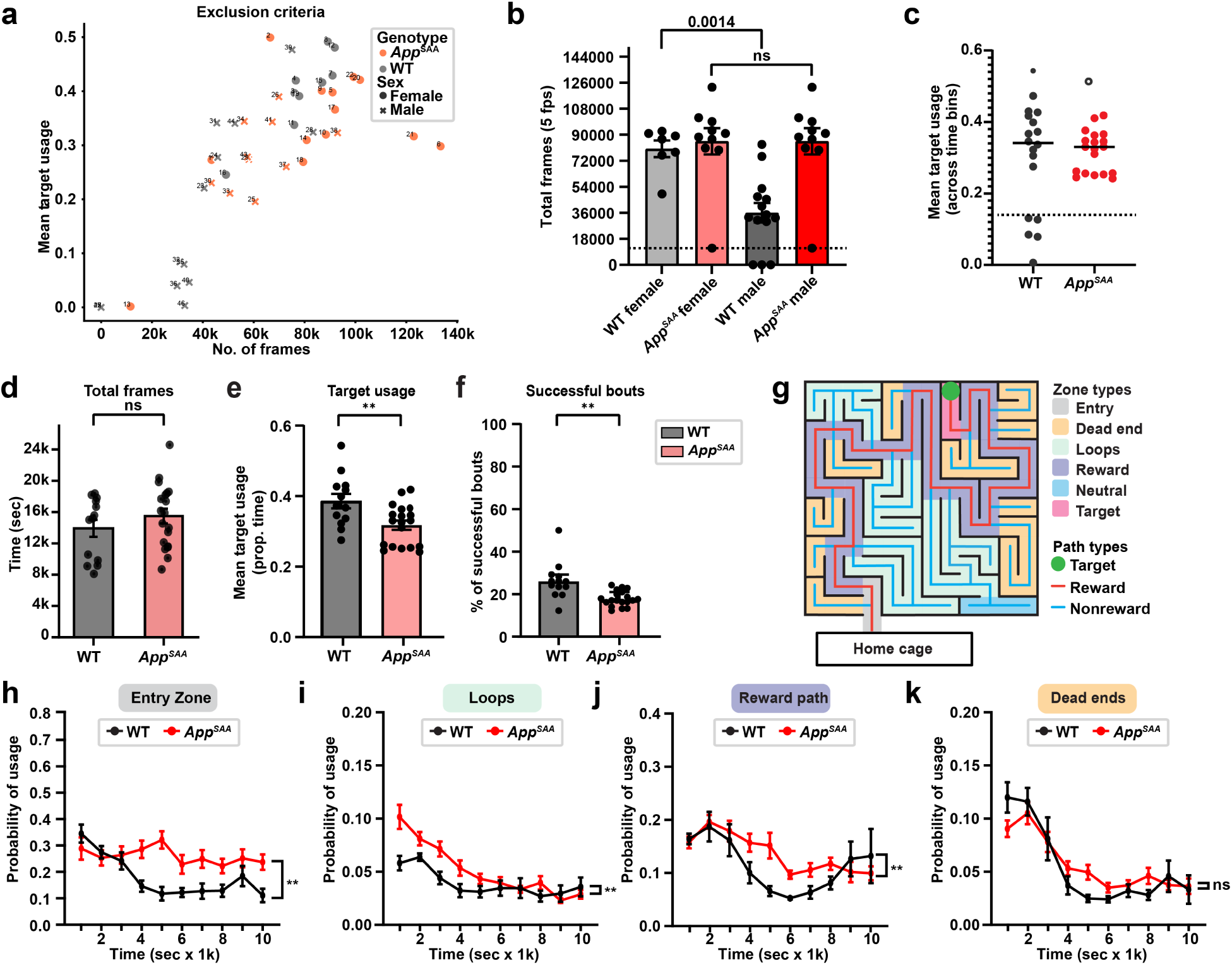
Impaired navigation to the target zone in *App*^SAA^ mice. 22-month-old *App*^SAA^ (*n*=19; 9F, 10M) and WT (*n*=13; 7F, 6M) mice were tested in the labyrinth. **a-c**, Exclusion criteria for selecting mice who performed the task from a cohort of *App*^SAA^ and WT mice. **a**, Scatterplot showing the total frames and mean target usage for each mouse, labeled by genotype and sex. **b**, Total frames in the maze with cutoff. **c**, Mean target usage with cutoff. **d**, Total frames in the maze were not changed amongst genotypes. **e,** Mean target zone usage. **f,** Percentage of successful bouts were reduced in *App*^SAA^ mice. Mann Whitney U-Test, \*\**p*<0.01. **g**, Diagram showing node types (non-decision, decision, and corner) and path types (reward, non-reward). **h-k**, Compared to WT, *App*^SAA^ mice displayed significant increases in usage of the entry zone (**h**), loops (**i**), and reward path (**j**). Repeated measures two-way ANOVA, \**p*<0.05, \*\**p*<0.01. Lines represent median ± sem (**h-k**).

**Supplementary Figure 5.**
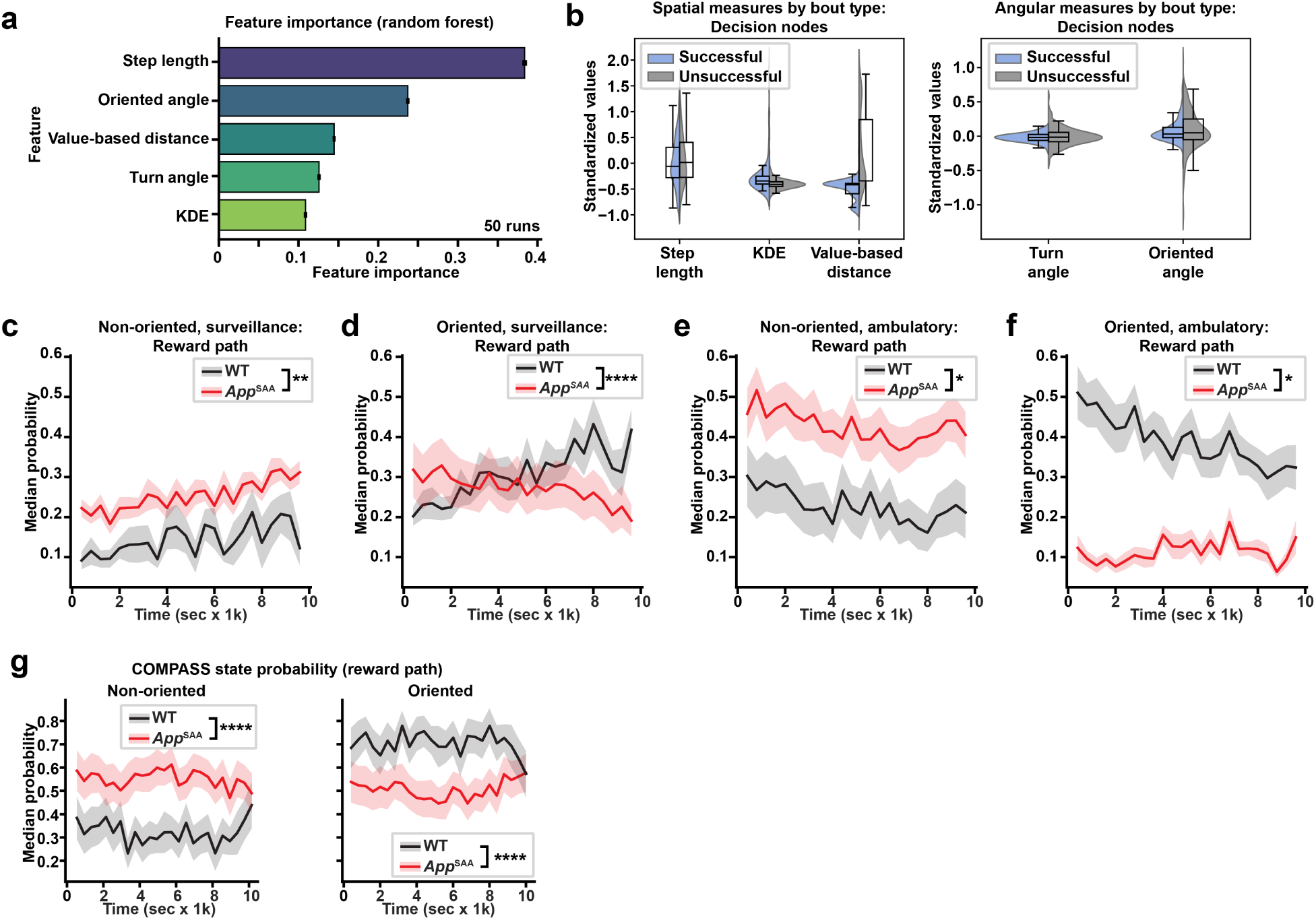
HHMM Model Validation and State Probability Along the Reward Path. 22-month-old *App*^SAA^ (*n*=19; 9F, 10M) and WT (*n*=13; 7F, 6M) mice were tested in the labyrinth. **a**, Feature importance computed using 50 Random Forest runs. **b**, Boxplots and distributions of spatial data streams (left) and angular data streams (right). **c-f**, Median probability of each CoMPASS state over time along the reward path. **g**, Median probability non-oriented (left) and oriented (right) states along the reward path for WT and *App*^SAA^ mice. Error bars represent median ± sem of each mouse. Repeated two-way ANOVA for probability x time. \**p*<0.05, \*\**p*<0.01, \*\*\**p*<0.001, \*\*\*\**p*<0.0001.

**Supplementary Figure 6.**
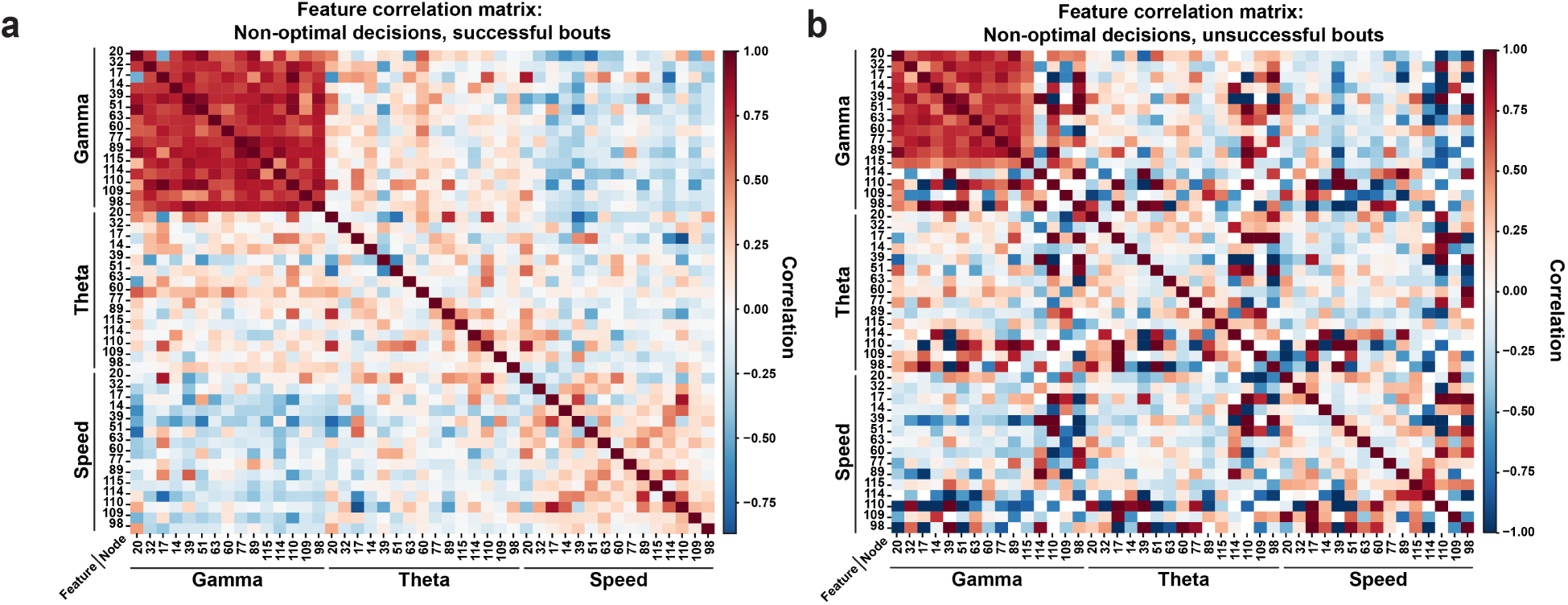
Non-stereotyped gamma power patterns during successful bouts. 5-mo-old WT mice (*n*=5, 5F) were implanted with wireless EEG/EMG probes in the PPC and recorded in the labyrinth maze during a 13 hr overnight trial. **a-b**, Correlation matrix heatmap between gamma, theta, and speed at decision nodes during non-optimal decisions (deviate from reward path) during successful (**a**) and unsuccessful (**b**) bouts.

**Supplementary Figure 7.**
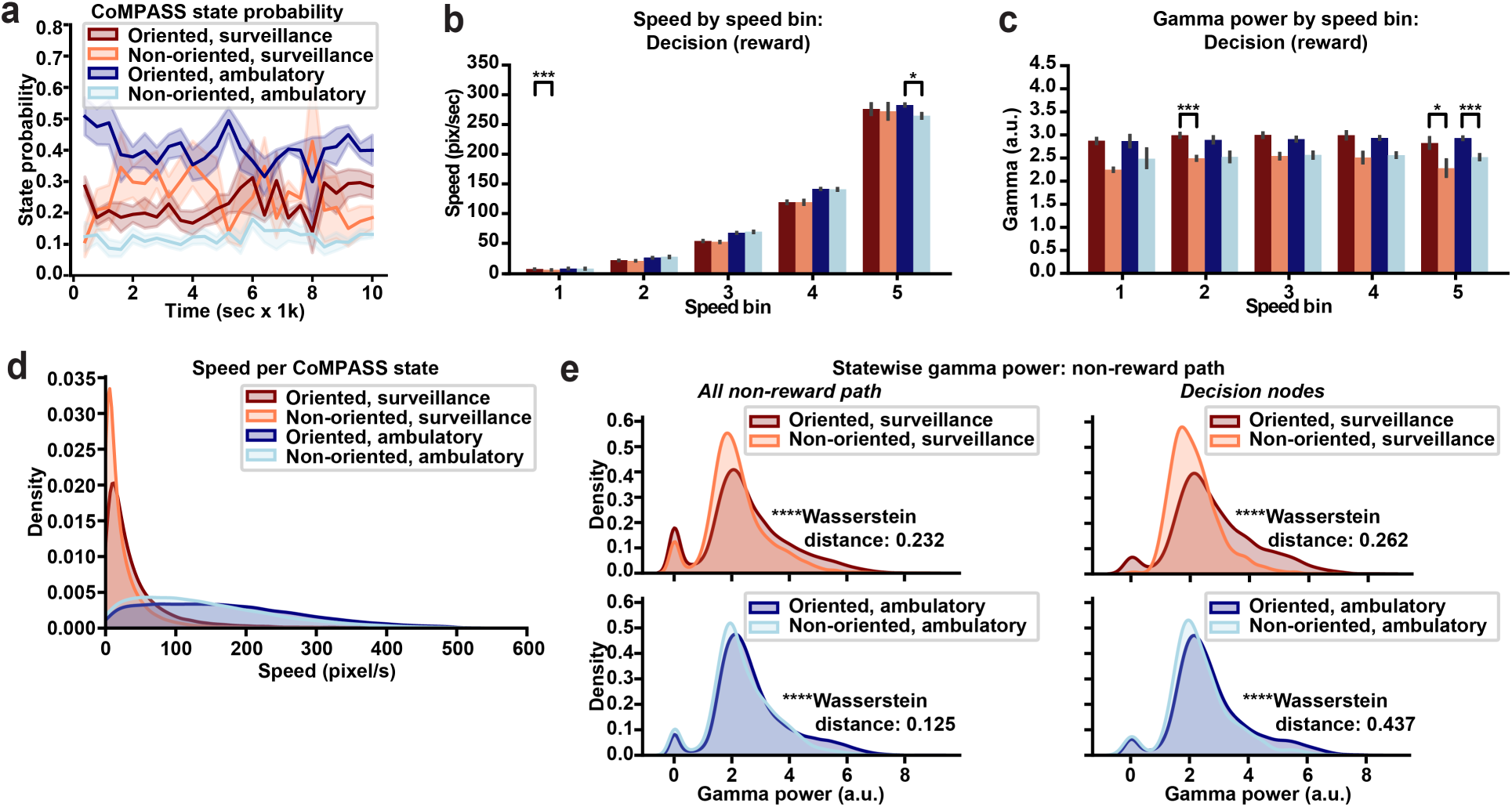
Increased gamma synchronization, independent of changes in speed. 5-mo-old WT mice (*n*=5, 5F) were implanted with wireless EEG/EMG probes in the PPC and recorded in the labyrinth maze during a 13 hr overnight trial. **a**, CoMPASS state probability in 400 second bins from 5-month-old wildtype (*n*=5; 5F) mice. Lines and shaded regions represent median ± sem. Statistics were obtained from a linear mixed effects model with mouse as a random factor. \**p*<0.05, \*\*\**p*<0.001. **b,c**, Bar plots the speed (**b**) and gamma power (**c**) for different bins of speed values. **d**, Distribution plots showing the speed per CoMPASS state. **e**, Distribution plots showing the normalized gamma power per CoMPASS state at all non-reward paths (left) or at decision nodes in non-reward paths (right). Statistics obtained from the Wasserstein distance. \*\*\*\**p*<0.0001.

## Supplementary Note

### Unique data streams for capturing goal-directed behaviors

To decode reward-oriented behavior in line with our definition, we generated complementary data streams. The oriented angle deviation captures the angular difference between the mouse’s current heading and the optimal trajectory toward the reward path. Thus, it captures the extent of goal-directed correction, where lower values indicate alignment with the reward path and higher values suggest deviation. By tracking this measure over time, we can infer whether reorientation efforts are directed toward exploiting learned pathways or represent exploratory deviations.

Second, the Value-based Distance metric is integral to assessing navigation efficiency. By combining egocentric and allocentric components, this measure incorporates the Euclidean distance to the target, which reflects immediate spatial proximity, and the grid-based path length, which accounts for the maze’s environmental complexity. This metric is essential for evaluating how efficiently the mouse navigates toward the goal. A lower value signifies direct, efficient movement, whereas a higher value indicates the need for additional reorientations or inefficient navigation strategies.

Lastly, Kernel Density Estimation (KDE) serves as a dynamic measure of spatial familiarity, reflecting the history of the mouse’s exploration and reinforcement of specific regions. KDE quantifies how often different maze areas are visited, allowing us to gauge the influence of past experiences on navigation. High KDE values suggest frequent exploration or learned preferences for certain paths, while lower values indicate less-utilized regions. This stream is vital for understanding whether reorientations toward the reward path are driven by learned associations or occur more randomly as part of an exploratory search.

Together, these data streams directly align with our objective of decoding reward orientation— defined here as the ability to reorient toward the reward path in pursuit of the target. The combination of real-time movement corrections (sternum reorientation), goal-directed navigation effort (value-based distance), and experience-driven spatial memory (KDE) provides a comprehensive framework for understanding how mice dynamically adjust their behavior to optimize reward acquisition.

### Leveraging Bayesian Gaussian Mixture Models (BGMM) and GMM-HMM for behavioral segmentation and temporal sequencing

To robustly segment behavioral states, we employed Bayesian Gaussian Mixture Models (BGMMs), which improve upon standard Gaussian Mixture Models (GMMs) by incorporating Bayesian priors to prevent overfitting and yield more stable state estimations. Unlike conventional clustering approaches, BGMMs utilize Dirichlet Process priors to determine the optimal clusters dynamically, allowing behavioral states to emerge organically rather than being arbitrarily predefined. To model the sequential dependencies, we employed Gaussian Mixture Model-Hidden Markov Models (GMM-HMMs), which incorporate Gaussian mixtures as emission distributions to accommodate variations within locomotor patterns that might otherwise be misclassified under discrete state assumptions. Furthermore, the Markovian framework allows for tracking behavioral evolution across bouts, offering insights into individual learning trajectories and adaptation processes.

By integrating BGMM for state segmentation with GMM-HMM for state transitions, we establish a hierarchical framework where locomotor states and other key data streams contribute to a feature space optimized for decoding reward orientation. This combination ensures the comprehensive modeling of both short-term movements and long-term navigational strategies. This structured pipeline ensures that every time point is simultaneously associated with a locomotor state and a reward-oriented state, allowing for a more granular and biologically meaningful characterization of navigation strategies. Unlike fixed-state HMMs, which impose rigid state definitions, BGMM-HMMs adaptively discover latent behavioral motifs, making them more suited for complex, real-world navigation tasks. Moreover, while reinforcement learning models such as Q-learning require predefined reward contingencies, our approach infers behavioral strategies in an unsupervised manner, enabling a more nuanced examination of genotype-specific adaptations in spatial learning and decision-making.

### Hierarchical multi-level framework for behavioral decoding

The hierarchical modeling approach refines locomotor states by embedding them within a task-relevant reward context. At Level 1, the decoding process categorizes behavior into Active Surveillance and Automated Ambulation, capturing fundamental movement patterns. However, these broad categories do not account for the decision-making processes that guide navigation within the task structure.

At Level 2, we introduce a higher-order classification by differentiating movement based on reward orientation. This step results in the emergence of two refined behavioral states:

1. Goal-Oriented Behavior – Movement patterns aligned with goal-directed navigation, where the mouse reorients toward or progresses along the reward path, in pursuit of the target zone.
2. Non-Goal-Oriented Behavior – Movements that do not contribute to immediate goal-directed progression, including deviations, pauses, or transitions into less optimal trajectories.

Since the hierarchical framework builds on previously decoded locomotor states, the resulting behavioral classification retains the structure of Active Surveillance and Automated Ambulation while incorporating reward context. This breakdown allows for points of overlap, where transitions between reward and non-reward states provide insight into decision-making dynamics. Thus, locomotor state sequences are further categorized as follows:

- Goal-Oriented + Active Surveillance
- Non-Goal-Oriented + Active Surveillance
- Goal-Oriented + Automated Ambulation
- Non-Goal-Oriented + Automated Ambulation

By combining unsupervised clustering, sequential state decoding, and hierarchical classification, this framework provides a fine-grained understanding of how mice modulate movement strategies in response to reward contingencies. The resulting state sequences offer insights into navigation efficiency, decision-making strategies, and cognitive flexibility, revealing how mice dynamically adjust their behavior to task demands.

## Methods

### Animals

The *App*^SAA^ knock-in mouse model (JAX stock #034711) was generated by introducing three Aβ humanization mutations—R684H, F681Y, and G676R—along with three familial Alzheimer’s disease (FAD) mutations: KM670/671NL (Swedish), E693G (Arctic), and T714I (Austrian) into the endogenous mouse *App* gene using a homologous recombination approach^29^. The mice are maintained on a C57BL/6J genetic background.

For the study, *App*^SAA^ knock-in mice and littermate wild-type (WT) control mice, including both sexes, were used. Mice were group-housed with access to water and food *ad libitum* and a standard 12-hour light-dark cycle. 18-22-month-old *App*^SAA^ mice (*n*=19; 9 females and 10 males) and WT controls (*n*=13; 7 females and 6 males) were used for the labyrinth experiments. All mouse experimental procedures were approved by the University of California, San Francisco Institutional Animal Care and Use Committee and were conducted in agreement with the NIH animal use guidelines.

### Maze design

Six custom mazes were designed with each measuring 18 (width) x 18 (length) x 1.5 (height) inches using opaque polycarbonate. 1.5-inch opaque polycarbonate walls were cut and glued to an 18 x 18-inch opaque polycarbonate ceiling, forming a unique design with loops, dead ends, and a target zone that can hold sucrose pellets. A 72 (width) x 36 (width) x 72 (height) inch metro rack was cut to have six 18-inch square cutouts. A clear acrylic base (0.5 inch thick) was fit to sit on two separate levels of the metro rack. The custom-built mazes fit in slots directly above the six 18-inch square cutouts. A 1-inch diameter hole was cut into the wall near the entry zone, through which a 1-inch acrylic tube connected to a home cage. The home cage contained bedding, nestling, a water bottle, and food *ad libitum*.

### Video acquisition

Six Basler acA1300-60gmNIR GigE cameras with 6mm CFF VIS-NIR lenses were mounted below the surface of the of the 18-inch square cutouts. The cameras were connected to two separate 4-port ethernet cards across two acquisition computers. Noldus Ethovision XT 17.0 was used to acquire the videos of resolution 1280 x 1024 with a frame rate of 5 fps. Because trials occurred during the animal’s dark cycle, multiple infrared illuminators were placed below the surface of the maze at an angle to reduce bright spots that occluded the mice.

### Experimental details

Six mice per trial were brought into the experimental room 30 minutes prior to the start of the experiment. One overnight 13hr trial was conducted starting at 6:30 pm and ending at 7:30 am, during the mice’s natural dark cycle. 20 sucrose pellets were placed in the shelf in the target zone and 1 pellet was placed at the entry zone. Mice were placed into their home cage at the start of the trial and given a water bottle and food ad libitum in their home cage. Mice had freedom to navigate between their home cage and the maze to search for sucrose pellets.

### Pose estimation

We used DeepLabCut^50^ to track 5 body parts. 50 frames were extracted from 12 videos, and 6 body parts parts (Nose, Left Flank, Right Flank, Sternum, Belly, and Tailbase) were manually labeled. A DeepLabCut (version 2.2.3) ResNet50 model was trained to 10^6^ iterations, resulting in train and test iterations of 0.91 pixels and 14.2 pixels, respectively. All videos used in this paper were analyzed using this pre-trained DeepLabCut network.

### Allocentric coordinate preprocessing, grid extraction, and bout definition

To determine the mouse’s specific location within the maze, we divided the maze into individual 12 x 12 (144 total) equal-sized individual grid cells. To account for any inconsistencies in DeepLabCut pose estimations, the x-,y-coordinates from all DeepLabCut keypoints were fed into our custom preprocessing script, which uses all possible grid cell transitions to determine if impossible jumps happened. These frames were removed from the downstream analyses. Using the pre-processed coordinates, the position was identified in terms of the grid nodes 1-144 in the labyrinth. Two rolling medians of the estimated (by DeepLabCut) likelihood of the mouse’s sternum position at a given frame were calculated using the 7 frames upstream and the other using the 7 frames downstream of the current frame. If either of the two calculated rolling medians of the estimated likelihood at any frame was below 0.2 then the grid node at that position was assigned as missing. Initial guesses of the entry times into the maze were estimated based on the frames with an assigned grid node position and no assigned grid position in the immediately preceding frame. Analogously, initial guesses of the exit times out of the maze were estimated based on the frames with an assigned grid node position and no assigned grid position in the immediate next frame. These entry and exit times define initial guesses of bouts into the labyrinth. These bouts were filtered so that the mean likelihood of the belly positions during each bout was greater than 0.8. Separate bouts that are less than or equal to 5 frames apart were merged. The entry times for each of these bouts were further updated to the first frame in the defined bout interval where the difference between the downstream rolling median likelihood and the upstream rolling median likelihood was greater than 0.8. Similarly, exit times for each of these bouts were further updated to the first frame in the defined bout interval where the difference between the upstream rolling median likelihood and the downstream rolling median likelihood was greater than 0.8. The lists of grid nodes visited during each bout that define the trajectory of the mouse during each bout were also recorded.

### Survival analyses

The path length or the number of (non-unique) nodes visited by the mouse before reaching a target zone for the first time since entering the maze during a bout was used as the response of interest. Two definitions of target zones were used here - one was the reward zone or grid nodes 84 and 85 where the sucrose pellets were placed and second was a neutral zone or grid nodes 107, 119, 131 and 143. Path lengths for a mouse in given bouts were considered censored if this mouse never reached the target zone and the path length for the entire bout (from entry to exit from the maze) was used in such situations. Per mouse, the distribution of the path lengths to the target zone for the first 20 visits were compared with the distribution of the path lengths for the subsequent 20 visits to the target zone using a survival analysis^51–53^. Specifically, Mixed Effects Cox Models (MECM) were used to model these data. The mouse ID was used as a random effect. MECMs were fit to estimate the hazard ratio comparing the path lengths between the first 20 visits to the target zone and the subsequent sets of 20 visits for the animals within each genotype (WT or *App*^SAA^) up to 120 visits in total and also between the genotypes for their 20-40 visits to the target zone. The idea is that hazard ratios greater than 1 would suggest, for example, that the mice are reaching the target zone using smaller path lengths in their subsequent 20-40 visits as compared to their path lengths during their first 0-20 visits. Hazard ratios less than 1 would suggest the opposite trend. The intuition is that if the mice were learning the path to the reward zone during their first 20 visits, the path lengths to reach the reward zone would be shorter for their subsequent 20-40 visits. Alternatively, if the mice learn that there is nothing of interest in their first 20 visits to the neutral zones then they would take longer or similar path lengths to reach the neutral zone during the subsequent 20-40 visits. Kaplan-Meier survival curves^51^ were plotted using the functionality in the survminer package^54^.

### Learning metrics

Within each bout, the number of frames spent on the reward path was divided by the number of frames in off-reward grid cells (deviation probability). An exponential decay function was fit to the deviation probability over time. For computing the probability of staying on the reward path at decision points, the number of occurrences were summed in which mice remained on the reward path within the next four frames after encountering a decision point and divided by the total number of occurrences.

### Binned heatmap usage and zone quantification

To create the regional proportion of usage heatmaps, we determined the total number of frames spent in the various regions during each 5,000 frame bin (1,000 sec) and normalized to the number of grid cells in each region. This only considered frames in which the mice were in the maze. Shannon’s entropy is used to measure the stochastic distribution of the proportion of usage across different regions. Here, we quantified the Shannon’s entropy for each mouse in bins of 5,000 frames (1,000 sec) for the duration of the trials. The total number of frames and proportion of target zone usage were used to exclude mice who did not perform the task from downstream analysis

### Simulated agent modeling

This analysis investigates optimal transitions in maze navigation, conditional on the mouse being at a Decision Node on the Reward Path at the current timestep (t). We define optimality as the extent to which mice follow the Reward Path in subsequent transitions. We consider two types of decision-making:

1. Single-Step Optimality – The transition is optimal if, at the next recorded spatial change (*t* + 1), the mouse remains on the Reward Path.
2. Two-Step Optimality – The transition is optimal if the mouse remains on the Reward Path for two consecutive spatial changes (*t* + 1 and *t* + 2).

We compare mice behavior to a random agent, which selects transitions at Decision Nodes based on uniform probability across all valid options.

#### Part 1: One-Step Optimality (Single-Step Decision Analysis) Defining One-Step Optimal Transitions

A transition is analyzed only if, at time *t*, the mouse is at a Decision Node located on the Reward Path. Given this condition, we evaluate whether the mouse remains on the Reward Path at the next spatial transition (*t* + 1).

Formally, Let:

- *G*_*t*_ be the grid occupied at *t*, where *G*_*t*_ is a Decision Node on the Reward Path.
- *G*_*t*+1_ be the next unique spatial location visited (i.e., the next grid where the mouse moves).

The transition is defined as optimal if:

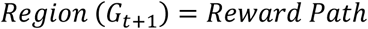

Where,

- *Region*(*G*) denotes the assigned region type of a given grid.

If the mouse transitions to any other region (e.g., a loop or dead-end), the transition is classified as non-optimal.

#### Preprocessing of Trajectory Data into Epochs

Each session’s trajectory data was first condensed to remove consecutive rows with the same Grid Number, retaining only unique grid positions (animal does not stay at the same grid node). This ensured that transitions reflected changes in spatial position rather than time spent in a single location. Each session-wise trajectory was segmented into clusters of 1000 frames. This segmentation allowed for evaluating performance trends across epochs. The total number of clusters *N*_*cluster*_ per session was calculated as:

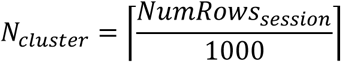

Where,

- *NumRows*_*session*_ is the total number of rows in a Session, or the total number of recorded time steps in a given session.

#### Identifying and Validating Grid Transitions

Transitions between grids were recorded when the mouse encountered a Decision Node. The validity of a transition was determined by checking whether the mouse moved from a given grid *G*_*t*_ to a valid next grid *G*_*t*+1_. Since the mouse’s movement speed and frame rate could result in skipping intermediate grid steps, a transition was considered valid if the next grid was within four steps from the current location in either direction. Let:

- *G*_*t*_ represent the current grid number at time step *t*
- *G*_*t*+1_ represent the next unique grid number at time step *t* + 1

The transition from *G*_*t*_ to *G*_*t*+1_ is valid if the mouse is moving from a Decision Node at *G*_*t*_. The set of valid transitions for a grid *G* is defined as:

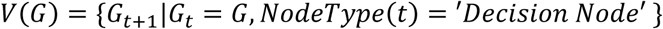

Where,

- *N*𝑜𝑑𝑒𝑇𝑦𝑝𝑒(*t*) is the type of node encountered at time step *t*

The optimal transition from a grid *G* is tracked if the subsequent grid *G*_*t*+1_ leads to the Reward Path. The set of optimal transitions 𝑂(*G*) is defined as:

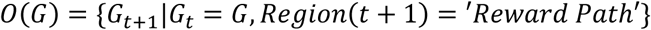

Where,

- *Region*(*t* + 1) represents the region the mouse moves into at time step t+1.

#### Evaluating Mouse Performance Against a Random Agent

The mouse’s decision-making performance was compared against a random agent, which simulated movement by randomly selecting a valid transition at each Decision Node. The performance metric at a grid *G* was defined as:

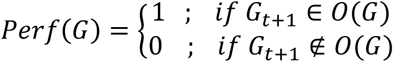

Where,

- 𝑃𝑒𝑟𝑓(*G*) = 1 if the chosen transition is optimal and 𝑃𝑒𝑟𝑓(*G*) = 0 otherwise.

For the random agent, the performance at grid *G* is averaged over 𝑛_𝑠𝑖*m*𝑢𝑙𝑎*t*𝑖𝑜𝑛𝑠_ trials, where each trial involved randomly selecting a next grid from the set of valid transitions and checking whether the transition is optimal. The random agent’s performance at grid *G* is given by the average over all the simulations:

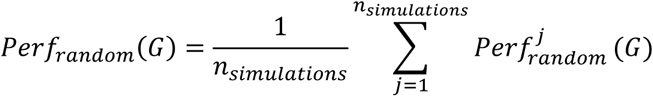

Where,

- 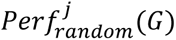 is 1 if the random agent’s j-th simulation resulted in an optimal transition, and 0 otherwise.

#### Bootstrap Confidence Intervals

To estimate the variability of performance metrics, bootstrap confidence intervals (CIs) were computed for both the mouse and random agent’s performance.

Let 𝑃𝑒𝑟𝑓 = [𝑃𝑒𝑟𝑓(*G*_1_), 𝑃𝑒𝑟𝑓(*G*_2_), …, 𝑃𝑒𝑟𝑓(*G*_𝑛_)] represent the performance

Where,

- 𝑛 is the number of decision nodes.

A bootstrap resampling method was applied to create a distribution of performance means:

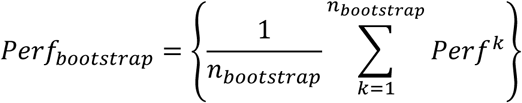

Where,

- 𝑃𝑒𝑟𝑓^𝑘^ is the performance in the k-th bootstrap resample.

The 95% confidence interval was estimated as:

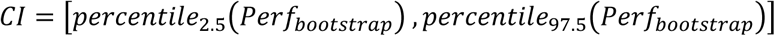

#### Relative Performance of the Mouse

A singular comparison metric, Relative Performance, was computed as the ratio of the mouse’s mean performance to that of the random agent:

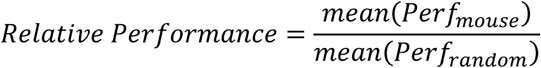

This metric quantifies the extent to which the mouse outperforms a random decision-making strategy, providing insight into the cognitive strategies underlying navigation behavior.

#### Chi-Square Test for One-Step Optimality

To evaluate whether mice exhibit decision-making behavior significantly different from chance levels, we applied a Chi-Square (𝜒^2^) test for independence to compare observed one-step optimal transition frequencies against expected frequencies from a random agent model. The Chi-Square test was conducted under the following hypotheses:

- Null Hypothesis (𝐻_0_): The probability of making an optimal transition (*t* → *t* + 1) is independent of whether the agent is a mouse or a random agent.
- Alternative Hypothesis ((𝐻_𝐴_): The probability of making an optimal transition is dependent on whether the agent is a mouse or a random agent, indicating a deviation from random behavior.

The Chi-Square statistic was computed as:

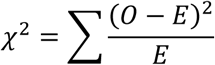

where:

- 𝑂 represents the observed frequency of optimal and non-optimal transitions for the mouse.
- 𝐸 represents the expected frequency of optimal and non-optimal transitions under the random agent model.

#### Significance Testing and Effect Size

A p-value was derived from the Chi-Square distribution with 1 degree of freedom, as we are comparing a categorical variable with two groups.

#### Exploration-Exploitation Model

The agent follows an exploitation-exploration decision strategy when transitioning from the current grid location *G*_*t*_ to the next grid *G*_*t*+1_. The transition probability is determined by a balance between exploratory and exploitative (non-exploratory) decision-making. The probability of selecting a transition is given by:

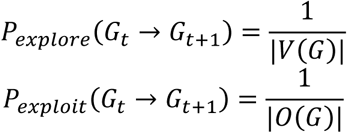

The agent makes decisions based on an exploration rate (ER), which determines the probability of selecting an exploratory vs. exploitative transition:

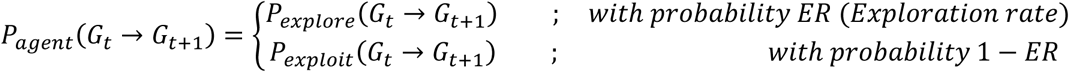

To evaluate the agent’s efficiency in reaching the reward path, performance is quantified as the proportion of transitions that lead to the reward path:

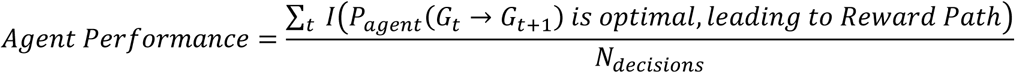

where:

- 𝐼(.) is an indicator function that returns 1 if the agent’s chosen transition is optimal (leading to the reward path) and 0 otherwise.
- *N*_𝑑𝑒𝑐𝑖𝑠𝑖𝑜𝑛𝑠_ is the total number of decisions made by the agent.

By systematically varying ER, we can analyze how different levels of exploration influence performance, allowing us to compare agent behavior across different conditions and genotypes.

### CoMPASS hierarchical modeling framework

#### Level 1: Locomotor Behavioral State,Inference

Understanding the latent structure of animal behavior requires probabilistic modeling techniques capable of inferring unobservable states from observed data. We employed a discrete-time Hidden Markov Model (HMM) to identify the underlying locomotor dynamics of mice navigating within the realms of a naturalistic setting. The HMM framework assumes that an observed time series *O* = (*O*_1_, *O*_2_, . . ., *O*_*T*_) is generated by a sequence of unobserved latent states *S* = (*S*_1_, *S*_2_, . . ., *S*_*T*_), where each state follows a Markov process with transition probabilities:

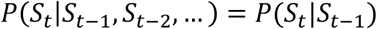

This first-order Markov assumption ensures that the probability of a state at time *t* depends only on the state at *t* − 1, making the model computationally tractable. The emission probabilities 𝑃(*O*_*t*_|*S*_*t*_) define how observable features (e.g., movement parameters) are generated given a hidden state.

#### Kinematic Feature Representation and State Transition Dynamics

Observed movement characteristics were derived from high-resolution positional tracking data (𝑥_*t*_, 𝑦_*t*_) sampled at 5 Hz. Two fundamental kinematic features—step length and turning angle— were computed for each timestep:

1. Step length (Δ𝑑_*t*_): Defined as the Euclidean distance between successive time points,

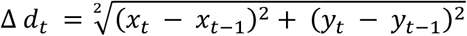

Step length values, constrained to be non-negative, exhibited a distribution resembling a gamma distribution, capturing the intrinsic variability of locomotor dynamics, where frequent short steps are interspersed with occasional longer strides, allowing for efficient navigation through the environment.

2. Turning (Bearing) angle (𝜃_*t*_): Defined as the angular deviation between successive movement vectors,

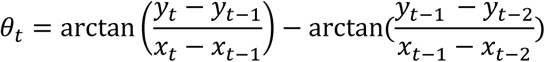

Turning angles followed a distribution resembling the von Mises distribution, which appropriately represents the cyclical nature of directional changes in locomotion. This distribution reflects the balance between smooth trajectory maintenance and abrupt reorientations, key components of adaptive navigation behavior.

#### Model Initialization and Parameter Estimation

To enhance model robustness and computational efficiency, we employed a grid search strategy for Level 1 parameter initialization^55^. Random samples for step length mean, standard deviation, turning angle mean, and concentration parameters were drawn from biologically plausible ranges. Model parameters were optimized by maximizing the likelihood function for the entire session and considering all sessions, iteratively updated using the Baum-Welch algorithm, a form of the Expectation-Maximization (EM) algorithm:

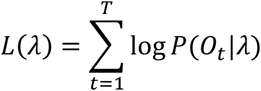

Where 𝜆 = (𝜋, 𝐴, 𝐵) is a vector of parameters consisting of the initial state probabilities 𝜋, transition matrix 𝐴, and the emission probabilities 𝐵.

#### Decoding Behavioral States

The most probable sequence of hidden states was inferred using the Viterbi algorithm, which finds the optimal state sequence *O*^∗^ that maximizes the posterior probability given the observations:

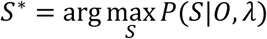

This decoding process revealed two primary locomotor states:

1. State 1: Active Surveillance—Characterized by low step lengths and high turning angles.
2. State 2: Automated Ambulation—Characterized by higher step lengths and lower turning angles.

#### Level 2: Reward-Guided Behavioral State Inference

While the HMM framework effectively captures locomotor states, identifying goal-directed behavior requires incorporating task-relevant features that provide critical context for spatial decision-making and reward proximity. By integrating these features, Level 2 modeling extends beyond movement patterns to infer behavioral states with greater specificity, distinguishing between reward-driven navigation and non-reward-oriented movement.

#### Feature Augmentation and Contextual Embedding

To infer reward-oriented behavioral states, we incorporated key spatial and decision-making features:

1. Oriented Angle Deviation (Δ𝜃): Oriented angle deviation quantifies the angular misalignment between the mouse’s current orientation and the optimal trajectory leading to the reward path. It is defined as the absolute difference between the heading direction vector of the mouse and the reference vector pointing toward the nearest reward-associated node. This measure captures the degree of goal-directed correction, where lower values indicate alignment with the reward path and higher values reflect deviation. The oriented angle deviation at time *t* is given by:

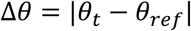

where,

- 𝜃_*t*_ is the mouse’s current heading direction, given by:

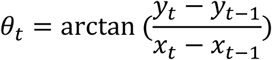

- 𝜃_𝑟𝑒𝑓_ is the optimal orientation towards the nearest node that ultimately leads to the reward path, computed as:

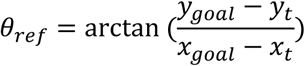

where (𝑥_*goal*_, 𝑦_*goal*_) is the nearest reference node leading to the reward path Since angles are inherently circular, we constrain Δ𝜃_*t*_ within the interval [−𝜋, 𝜋], ensuring a properly scaled measure of deviation. By tracking Δ 𝜃_*t*_ over time, we can infer reorientation behavior, distinguishing between goal-directed corrections and exploratory deviations in spatial navigation.

2. Value-Based Distance (𝑑_𝑣𝑎𝑙𝑢𝑒_): To capture the relationship between distance and navigation complexity, we computed a value-based distance metric, defined as the product of Euclidean displacement to the target (𝑑_𝐸𝑢𝑐𝑙𝑖𝑑𝑒𝑎𝑛_) the grid-based path length (navigation cost) required to reach the goal (*N*_*g*𝑟𝑖𝑑𝑠_ ∈ [1, 55]). This metric integrates both spatial proximity and path complexity, providing a more complete measure of goal-directed effort.

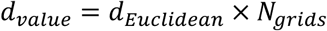

where 𝑑_𝐸𝑢𝑐𝑙𝑖𝑑𝑒𝑎𝑛_ is the Euclidean distance from the current position to the target and *N*_*g*𝑟𝑖𝑑𝑠_ is the number of traversed grid cells.

3. Kernel Density Estimation (KDE): To quantify spatial occupancy patterns, we applied Kernel Density Estimation^56^ to the mouse’s positional data over time. The optimal bandwidth was determined using a grid search with 5-fold cross-validation, ensuring robustness and generalizability across sessions.

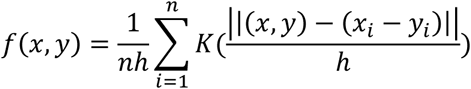

where 𝐾(⋅) is the Gaussian kernel, and ℎ is the bandwidth.

The Level 1 decoded states (Active Surveillance, Automated Ambulation) are also integrated into the feature set, providing a hierarchical behavioral representation.

#### Modeling Framework and Initialization

To classify fine-scale behaviors, we employ a hierarchical statistical model composed of:

1. Bayesian Gaussian Mixture Model (BGMM) for adaptive clustering of behavioral features.
2. Gaussian Mixture Model Hidden Markov Model (GMM-HMM) for state sequence decoding with temporal dependencies.

#### Bayesian Gaussian Mixture Model (BGMM) Initialization

To identify distinct behavioral clusters in an unsupervised manner, we fit a BGMM with Dirichlet Process priors^57^. The probability of an observation *O* given the mixture model is:

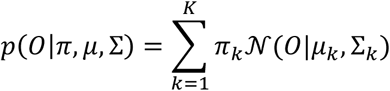

Where,

- 𝐾 represents the number of Gaussian components (clusters) in the mixture model.
- 𝒩(𝜇_𝑘_, Σ_𝑘_) is the Gaussian component with mean 𝜇_𝑘_ and covariance Σ_𝑘_
- 𝜋_𝑘_ is the mixing weight, drawn from a Dirichlet prior

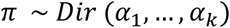

where the concentration parameter 𝛼 controls component sparsity. This approach prevents overfitting and allows adaptive determination of behaviorally relevant clusters.

The Dirichlet Process prior allows the model to determine the number of clusters adaptively, preventing overfitting and ensuring generalization across different experimental sessions. To prevent numerical instability, covariance regularization is applied. The learned cluster parameters from the BGMM serve as initialization values for the subsequent GMM-HMM, which further refines the latent state sequence by integrating temporal dependencies.

#### Gaussian Mixture Hidden Markov Model (GMM-HMM) for Sequential State Decoding

The GMM-HMM extends the clustering output from the BGMM by introducing latent state transitions, allowing us to capture the temporal structure of behavioral sequences. This probabilistic framework models the observed behavioral features as emissions from hidden states that evolve over time according to a first-order Markov process^58^. For a set of discrete latent states that correspond to distinct behavioral modes, the emission probability of an observation *O*_*t*_ given the latent state *S*_*t*_, is modeled as a Gaussian mixture distribution:

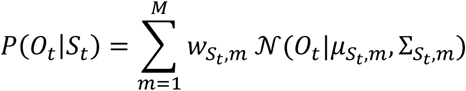

Where,

- 𝑀 represents the number of mixture components
- 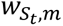 is the weight of component *m* in state *S_t_*
- 𝒩(*O*_*t*_|𝜇_*S_t_*,*m*_, Σ_*S_t_*,*m*_) denotes a Gaussian distribution with mean 𝜇 and covariance Σ.

The GMM-HMM is trained using the Expectation-Maximization (EM) algorithm with the Baum-Welch method^59^, which iteratively estimates the transition probabilities (𝐴_𝑖𝑗_ = 𝑃(*S*_*t*+1_ = *j*|*S*_*t*_ = 𝑖)) of the 𝐾 × 𝐾 (for 𝐾 hidden states) transition matrix 𝐴^𝐾×𝐾^ and emission distributions (𝒩(𝜇, Σ)) until model convergence. To prevent overfitting and ensure optimal convergence, we apply early stopping with a patience window.

Early stopping monitors the model’s negative log-likelihood (NLL) on a validation set, halting training when performance ceases to improve over a specified number of iterations (𝑃). If the improvement in NLL falls below a predefined threshold within the patience window, the optimization terminates early, selecting the best-performing model parameters before overfitting occurs^55^. Formally, training halts when:

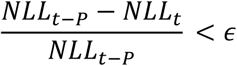

where *t* is the current iteration, 𝑃 is the patience window, and 𝜖 is the improvement threshold. This approach ensures robust generalization across sessions while reducing unnecessary computational expense. Once trained, the optimal latent state sequence is inferred using the Viterbi algorithm, which computes the most likely state path given the estimated transition and emission probabilities. This step ensures a globally optimal assignment of behavioral states, minimizing spurious fluctuations in classification.

#### Model selection, validation & interpretation

The final model was derived through a structured process prioritizing both statistical generalization and behavioral interpretability—tailored for decoding latent navigational strategies.

##### 1. Hyperparameter Optimization

Model construction began with a systematic grid search across key hyperparameters governing both the BGMM and the GMM-HMM, primarily:

- Number of latent states,
- Number of Gaussian mixtures per state,
- Covariance regularization strength,
- Dirichlet prior concentration to control mixture sparsity.

To ensure numerical stability and prevent overfitting, full covariance matrices from the BGMM were regularized. These parameter combinations were tested iteratively across multiple model instantiations.

##### 2. Phase-Aligned Cross-Session Model Evaluation

To rigorously evaluate generalizability across sessions and time, we applied a phase-aligned leave-one-session-out cross-validation framework:

- Each session was divided into equal-length behavioral epochs (e.g., early to late navigation).
- At each fold, one session-phase pair was held out, and the remaining sessions (same phase) were used for training/validation.

This preserved temporal alignment of behavioral progression while testing the model’s ability to generalize across individuals. Each model was evaluated on held-out data using:

- Log-Likelihood
- Akaike Information Criterion (𝐴𝐼𝐶)^60^

##### 3. Latent State Decoding and Behavioral Interpretation

Following selection of the optimal model, the decoded latent states were interpreted post hoc using a heuristic, data-driven approach. Rather than applying rigid, predefined decoding rules, we employed qualitative thresholding guided by the structure of the input data. Each state’s identity was inferred based on internal consistency across features, and alignment with the theoretical behavioral structure.

### Wireless EEG transmitter implantation and telemetry recordings

HD-X02 mouse transmitters (Data Sciences International) was performed according to the manufacturer’s surgical protocol for EEG and neck EMG in mice. Mice were anesthetized with Isoflurane (3% induction, 1% maintenance) and administered buprenorphine (0.05 mg/kg) prior to surgery. Ketoprofen (5 mg/kg) was administered at the end of surgery for analgesia. A large incision was made posterior of the eyes towards the cerebellum, and the transmitter was implanted into a subcutaneous pocket along the left dorsal flank. Two bipolar EMG leads were placed into the cervical trapezius muscles in the dorsal region of the neck, and two EEG leads were placed over the Posterior Parietal Cortex (+1.6 AP, -1.9 AP, 0 DV) and the Cerebellum. Transmitter batteries were turned on prior to the labyrinth recording, and the Ponemah software was used to acquire EEG and EMG. EEG and EMG were collected at 500 Hz. The Raw signal collected TTL pulses from Noldus Ethovision XT to align the behavior to the EEG data.

### EEG analysis

Recordings from the Ponemah software were converted into edf files that were then analyzed using custom Python code. Movement artifacts or drops in the signal were first found by z-scoring the signal and finding points where –6 > z_score > 6. A linear interpolation was used to remove these points. We computed a multitaper PSD (MNE-Python^61^) from this signal in 1 s time bins with a sliding window of 0.2 seconds to estimate the power in 1 Hz frequency bins. We then took the integral between our frequency bands 5-12 Hz (theta) and 27-100 Hz (gamma) to obtain our power measurements. This sliding window method allowed us to obtain power measurements for each frame of the video. The raw powers were normalized to the 10^th^ percentile of times when the mice were in the maze.

The normalized powers were then used to compute the mean power per Level 1 CoMPASS state at nodes along the reward path, during all bouts and split by successful versus unsuccessful bouts. The feature correlation matrices were created by taking the normalized theta powers, gamma powers, and speed when mice make correct or incorrect decisions at decision nodes during successful or unsuccessful bouts. The mean correlation is computed across all events and mice. Histograms of the gamma powers were also created for when mice are operating in the different CoMPASS states at different node and path types. The Wasserstein distance of the distributions was computed to compare statistical significance.

### Supervised classification of bout success

To evaluate the ability of behavioral and neural features to predict successful navigation, we employed a phase-based cross-validation framework across sessions.

#### Phase-Based Segmentation

Each session was divided into temporally ordered bins of 2000 frames, assigning a phase label ϕ ∈ {1, 2, …, Nϕ} as:

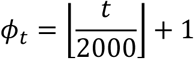

For,

- Frame *t* in a Session

#### Cross-Validation Framework

For each phase ϕ, we define:

- *S_ϕ_* = {*s_1_*, . . ., *s_k_*}: all sessions within phase ϕ,
- *D_ϕ_* = ⋃ *D_s_* for all *s* in *S_ϕ_* : the data pooled from those sessions.

For every session *s_i_* ∈ *S*_ϕ_, we trained on 𝐷_𝜙_ \ 𝐷_𝑠𝑖_ and evaluated on 𝐷ₛ_ᵢ_:

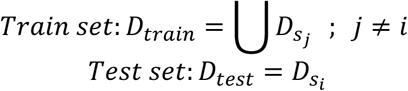

#### Feature Engineering

Let X ∈ ℝⁿˣᵈ denote the feature matrix with d behavioral and neural predictors (e.g., step length, turn angle, gamma/theta power etc.), and y ∈ {0,1}ⁿ the binary outcome indicating whether a bout was successful (i.e., the animal reached the target zone). All features were standardized via z-score normalization using parameters fit on the training data: ẋᵢⱼ = (xᵢⱼ − μⱼ) / σⱼ, where μⱼ and σⱼ are the mean and standard deviation of feature j in the training fold

#### Modelling Framework & Evaluation Metrics

We benchmarked three widely-used classification models:

- Logistic Regression62 with class balancing and L2 regularization
- Random Forests63, an ensemble of decision trees trained on bootstrapped samples
- XGBoost^64^, an efficient gradient-boosted decision tree model.

Each model was trained on the training folds and evaluated on the held-out session within the phase. Predicted probabilities were recorded for each test set. Model performance was assessed using ROC AUC, precision-recall AUC, and fold-wise class distributions. Feature attribution for the XGBoost model was computed using SHAP (SHapley Additive exPlanations) values^65^.

### UMAP embedding of gamma oscillatory dynamics during decision-making

To capture the temporal structure of gamma oscillatory power and its relation to latent behavioral states, we employed a dimensionality reduction strategy using Uniform Manifold Approximation and Projection (UMAP) on gamma power segments centered around key decision nodes.

#### Sliding-Window Feature Construction

Gamma power traces (𝛾(*t*)) were first segmented using a fixed-length sliding window approach:

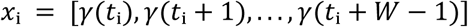

where 𝑊 = 20 is the window size and 𝑥ᵢ ∈ 𝑅²⁰ represents the gamma power segment for the i-th window. Only windows for which the middle frame corresponds to a pre-defined decision node were retained. All gamma values were z-scored before embedding.

#### UMAP Embedding

We applied UMAP^66^, a non-linear dimensionality reduction technique, to reduce the windowed gamma features into a low-dimensional manifold. UMAP preserves both local and global structure by minimizing the cross-entropy between high-dimensional and low-dimensional fuzzy simplicial sets. The following UMAP parameters were used:

- n_neighbors = 15
- min_dist = 0.1
- metric = ‘euclidean’
- n_components = 3

## Data Availability

All data used in this study are publicly available upon publication.

## Code Availability

The CoMPASS modeling framework and labyrinth analysis pipeline are available at https://github.com/CoMPASS-Framework/CoMPASS-Labyrinth. This repository contains example data and scripts for generating the results.

## Acknowledgements

We thank Pascal Sanchez and Dan Xia (Denali Therapeutics) for the *App*^SAA^ mice. We thank the Palop Lab members and LARC staff for assistance with mouse breeding, husbandry, and further technical support. The study was supported by US National Institutes of Health (NIH) grants RF1AG062234, R01AG062629, and P01AG073082 (J.J.P.).

## Author Contributions

P.S.H., S.C.B., and J.J.P. conceived the project and interpreted all data. P.S.H performed the animal surgeries and experimentation. P.S.H. and J.J.P. designed the labyrinth maze. S.C.B. designed CoMPASS and the models. P.S.H. and S.C.B. performed data analyses. R.R.T. contributed to survival analysis and statistical analysis. N.K. performed pose estimation. D.X. provided mice. P.S.H., S.C.B., and J.J.P. wrote the manuscript with contributions from all authors. J.J.P. supervised the project.

## Competing Interests

D.X. works for Denali Therapeutics. The other authors declare no competing interests.

